# The primary cilium controls programmed cell death via its proteasome-regulating function

**DOI:** 10.64898/2026.01.07.698216

**Authors:** Antonia Wiegering, Sandra Hägele, Stefanie Kuschel, Monika Böddeker, Christiane Ott, Sophie Saunier, Tilman Grune, Ulrich Rüther, Sylvie Schneider-Maunoury, Peter Walentek, Thorsten Pfirrmann, Christoph Gerhardt

## Abstract

Primary cilia are tiny cellular protrusions of nearly every vertebrate cell controlling multiple cellular processes, such as proliferation, differentiation, migration etc. Their dysfunction results in severe human diseases collectively referred to as ciliopathies. Remarkably, many ciliopathies are associated with increased programmed cell death (PCD). However, it is largely unknown how primary cilia regulate PCD. In *in vitro* (murine and human cells) and *in vivo* (*Xenopus laevis* and mouse) models, we observed elevated PCD in the absence of the ciliopathy protein RPGRIP1L. Mechanistically, our data elucidated that RPGRIP1L controls PCD by governing the activity of the ciliary proteasome. By using super-resolution microscopy, we first showed that the apoptosis inducer MOAP1 localises to primary cilia. Furthermore, our investigations revealed that RPGRIP1L controls PCD via the degradation of MOAP1 by the ciliary proteasome. Based on our finding that two more ciliopathy proteins, TCTN1 and CEP290, modulate PCD via regulating the activity of the ciliary proteasome, we suggest that the proteasomal degradation of MOAP1 represents a general mechanism by which primary cilia control PCD.

## Introduction

Ciliopathies are severe, often deadly human diseases caused by the dysfunction of tiny cell protrusions known as primary cilia. Once considered to be rare diseases, the number of ciliopathies is permanently rising ^1^. Even common diseases such as cancer and neurodegenerative disorders are among the diseases which are associated with defective primary cilia ^2–7^. Basically, the primary cilium consists of three different parts – the basal body (BB), the transition zone (TZ) and the axoneme. As a modified mother centriole, the BB is the origin of the ciliary microtubule-based axoneme. The axoneme, formed by nine doublet microtubules arranged in a circle, provides stability to the cilium and is essential for the transport of proteins that are conveyed through the cilium. A region of about 0.5 μm, present at the proximal end of the axoneme and designated as the TZ, functions as the ciliary gatekeeper, controlling ciliary protein import and export (ciliary gating) and hence ciliary protein composition ^1^.

Primary cilia function as signalling hubs and, as such, they are deeply involved in the regulation of cellular processes such as proliferation, differentiation, migration etc. ^8^. Remarkably, the truncation or complete loss of several ciliary proteins lead to increased cell death suggesting an important role of primary cilia in the regulation of PCD (summarised in ^9^). In line with these studies, PCD has been described as a pathological feature of polycystic kidney disease (PKD), a renal ciliopathy ^10,11^. As many ciliopathies can be traced back to mutations in genes encoding TZ proteins ^1^, it is remarkable that mutations in genes encoding TZ proteins provoke enhanced apoptosis in rats and mice. For example, Transmembrane Protein 67 (TMEM67 alias MKS3) deficiency causes an increase of apoptosis in rat eyes and in the murine cerebellum ^12,13^. The loss of Tectonic-2 (TCTN2) increases apoptosis in murine ventral neuroectoderm and facial ectoderm, while the absence of Tectonic-3 (TCTN3) leads to enhanced apoptosis in the brain of mouse embryos ^14,15^. Although numerous studies point to a regulation of PCD by primary cilia, the mechanisms by which primary cilia control PCD are poorly understood and may be indirect and cell type-specific. Here, we describe a novel mechanism underlying the cilia-mediated control of PCD.

By using both *in vitro* (murine and human cell lines) and *in vivo* (*Xenopus laevis* and mice) models with targeted inactivation of distinct TZ proteins, combined with super-resolution imaging approaches [Airyscanning and direct stochastic optical reconstruction microscopy (dSTORM)], subcellular proteasomal activity assays, CoIP analysis, proximity ligation assays as well as small molecule- and RNAi-based rescue experiments, we reveal that TZ proteins orchestrate the regulation of the ciliary proteasome which, in turn, governs PCD by controlling the amount of the Modulator of Apoptosis 1 (MOAP1) at the ciliary base.

## Results

### PCD is increased by absence, deficiency or truncation of the TZ protein Retinitis Pigmentosa GTPase Regulator Interacting Protein 1 Like (RPGRIP1L)

We and others showed that mutations in the gene encoding the TZ protein RPGRIP1L result in various, very severe, often deadly ciliopathies ^16–18^. Since many ciliopathies are associated with increased PCD, we investigated PCD in mammalian *in vitro* models. *Rpgrip1l*^-/-^ NIH3T3 cells were serum-starved for 24 h to induce ciliogenesis and immunostained with antibodies against the PCD markers cleaved caspase-3 (CC3) and cleaved caspase-9 (CC9). Significantly more serum-deprived *Rpgrip1l*^-/-^ NIH3T3 cells were positive for CC3 and CC9 than wild-type NIH3T3 cells in the same conditions (Fig. 1A,B). As cell death and cell proliferation are opposing cellular processes determining the number of cells, we also quantified the number of proliferating cells in *Rpgrip1l*^-/-^ NIH3T3 cells by using an antibody against phospho-histone H3 (pH3). The number of pH3-positive *Rpgrip1l*^-/-^ NIH3T3 cells was unaltered (Fig. EV1A) indicating that the loss of RPGRIP1L does not affect proliferation in NIH3T3 cells. In order to test whether the increase in PCD by the loss of RPGRIP1L is cilia-dependent, we investigated PCD in *Rpgrip1l*^-/-^ NIH3T3 cells without serum deprivation. As expected, the percentage of ciliated cells was significantly higher in serum-deprived *Rpgrip1l*^-/-^ NIH3T3 cells than in *Rpgrip1l*^-/-^ NIH3T3 cells without serum starvation (Fig. 1C). Moreover, significantly less *Rpgrip1l*^-/-^ NIH3T3 cells without serum deprivation were CC3-positive than in the serum-deprived condition (Fig. 1D). In contrast, in wild-type NIH3T3 cells, while the percentage of ciliated cells was increased upon serum deprivation, no difference was observed in the number of CC3-positive cells, indicating that the degree of ciliation does not affect PCD in wild-type NIH3T3 cells. Of note, despite a lower ratio of ciliation, serum-starved *Rpgrip1l*^-/-^ NIH3T3 cells had a significantly higher ratio of CC3 positivity than wild-type cells in the same conditions (Fig. 1C,D). Taken together, these findings indicate that RPGRIP1L regulates PCD in NIH3T3 cells in a cilia-dependent manner.

**Fig. 1:**
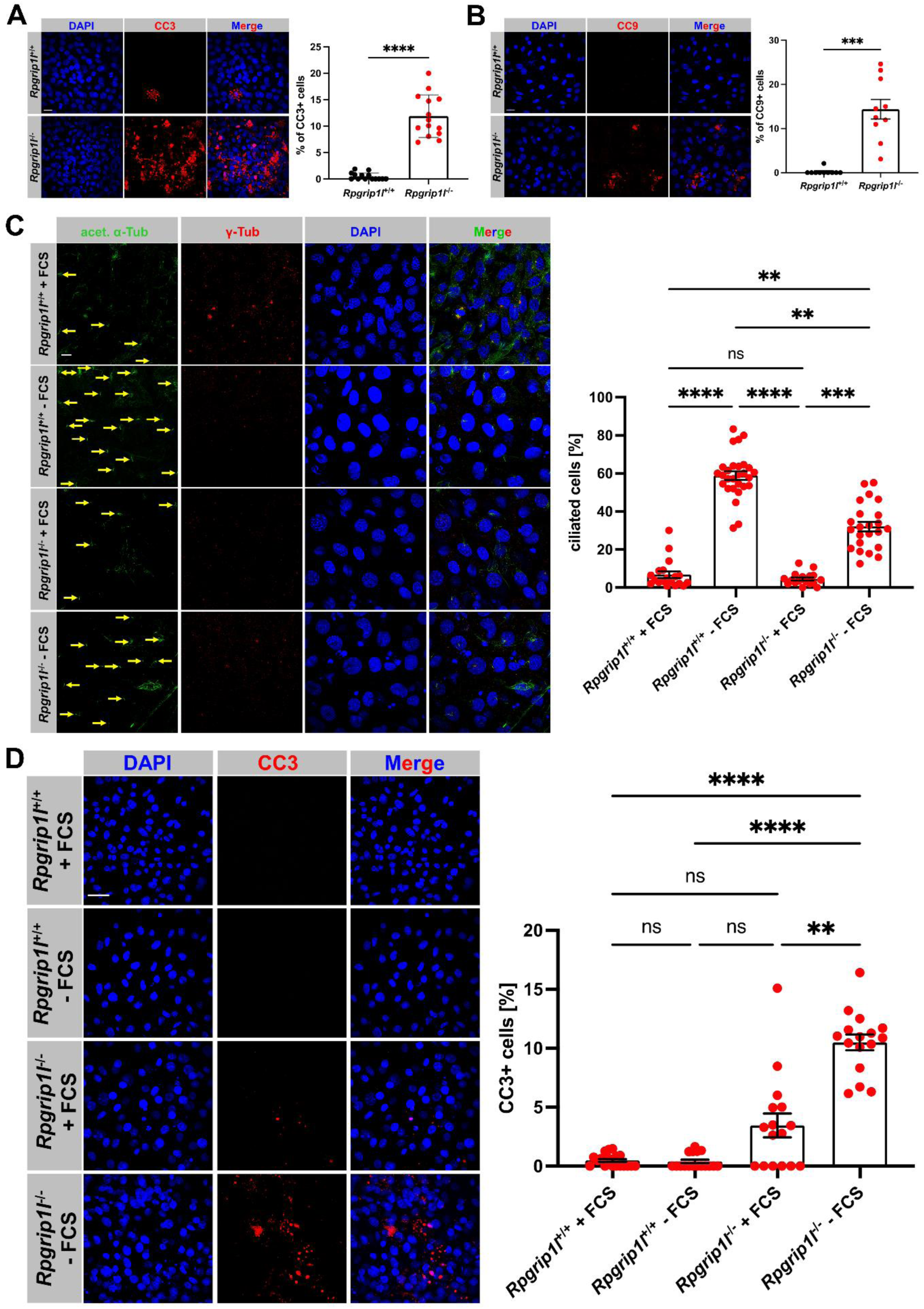
Loss of RPGRIP1L results in a cilia-dependent increase of PCD in NIH3T3 cells. (**A-D**) Immunofluorescence on NIH3T3 cells. Cell nuclei are stained in blue by DAPI. (**A**) Cells undergoing PCD are marked in red by CC3. Cells are serum-deprived. The scale bar (in white) represents a length of 20 μm. At least 1600 cells per genotype were used for these measurements, pooled from three replicates. Data shown as mean ± s.e.m. Statistical evaluation was performed by using two-tailed unpaired t test with Welch’s correction. Asterisks denote statistical significance (**** *P* <0.0001). (**B**) Cells undergoing PCD are marked in red by CC9. Cells are serum-starved. The scale bar (in white) represents a length of 20 μm. At least 450 cells per genotype were used for quantification, pooled from three replicates. Data shown as mean ± s.e.m. Statistical evaluation was performed by using two-tailed unpaired t test with Welch’s correction. Asterisks denote statistical significance (*** *P* = 0.0001). (**C**) The ciliary axoneme is stained in green by acetylated α-tubulin and the BB in blue by γ-tubulin. Yellow arrows point to cilia. Serum starvation is indicated by -FCS. The scale bar (in white) represents a length of 10 μm. At least 770 cells per genotype and treatment were used for quantification, pooled from three replicates. Data shown as mean ± s.e.m. Statistical evaluation was performed by using Kruskal-Wallis test with Dunn’s multiple comparisons test. Asterisks denote statistical significance (*Rpgrip1l*^+/+^ + FCS vs. *Rpgrip1l*^+/+^ - FCS: **** *P* <0.0001; *Rpgrip1l*^+/+^ + FCS vs. *Rpgrip1l*^-/-^ - FCS: ** *P* = 0.0014; *Rpgrip1l*^+/+^ - FCS vs. *Rpgrip1l*^- /-^ + FCS: **** *P* <0.0001; *Rpgrip1l*^+/+^ - FCS vs. *Rpgrip1l*^-/-^ - FCS: ** *P* = 0.0065; *Rpgrip1l*^- /-^ + FCS vs. *Rpgrip1l*^-/-^ - FCS: *** *P* = 0.0007). ns = not significant. **d,** Cells undergoing PCD are marked in red by CC3. Serum starvation is indicated by -FCS. The scale bar (in white) represents a length of 30 μm. At least 900 cells per genotype and treatment were used for these measurements, pooled from three replicates. Data shown as mean ± s.e.m. Statistical evaluation was performed by using Kruskal-Wallis test with Dunn’s multiple comparisons test. Asterisks denote statistical significance (*Rpgrip1l*^+/+^ + FCS vs. *Rpgrip1l*^-/-^ - FCS: **** *P* <0.0001; *Rpgrip1l*^+/+^ - FCS vs. *Rpgrip1l*^-/-^ - FCS: **** *P* <0.0001; *Rpgrip1l*^-/-^ + FCS vs. *Rpgrip1l*^- /-^ - FCS: ** *P* = 0.0037). ns = not significant (*Rpgrip1l*^+/+^ + FCS vs. *Rpgrip1l*^+/+^ - FCS: *P* >0.9999; *Rpgrip1l*^+/+^ + FCS vs. *Rpgrip1l*^-/-^ + FCS: *P* = 0.3974; *Rpgrip1l*^+/+^ - FCS vs. *Rpgrip1l*^- /-^ + FCS: *P* = 0.1449).

Recently, we demonstrated that RPGRIP1L deficiency in mouse spinal organoids results in defects that differ from those observed in human spinal organoids ^19^. To investigate whether RPGRIP1L is also able to regulate PCD in human cells, we analysed its role in the regulation of PCD in the HEK293 (human embryonic kidney 293) cell line. We found that PCD was enhanced in serum-deprived *RPGRIP1L*^-/-^ HEK293 cells, compared to wild-type cells (Fig. 2A). Moreover, PCD was rescued by re-expressing full-length RPGRIP1L in serum-deprived *RPGRIP1L*^-/-^ HEK293 cells (Fig. 2A). Most ciliopathy patients suffering from mutations in *RPGRIP1L* do not lack the entire protein but produce a truncated or slightly modified form of RPGRIP1L. So, we tested whether a ciliopathy patient variant affects PCD. Previously, it was reported that a mutation in *RPGRIP1L* which results in the amino acid alteration T615P represents the most common mutation in *RPGRIP1L* causing disease in about 8-10% of Joubert syndrome type B patients negative for *nephrocystin 1* (*Nphp1*), *centrosomal protein 290* (*Cep290*) or *Abelson helper integration site 1* (*Ahi1*) mutations ^18^. Consequently, we transiently transfected serum-deprived *RPGRIP1L*-negative HEK293 cells with a plasmid encoding the T615P mutation (referred to as RPGRIP1L-T615P ^16^). In comparison to serum-deprived wild-type HEK293 cells, PCD was significantly increased in serum-deprived *RPGRIP1L*-negative HEK293 cells transfected with RPGRIP1L-T615P (Fig. 2A). Interestingly, there was no difference in the number of CC3-positive cells between serum-deprived *RPGRIP1L*-negative HEK293 and serum-deprived *RPGRIP1L*-negative HEK293 cells transfected with RPGRIP1L-T615P (Fig. 2A). As seen in NIH3T3 cells, proliferation was not altered in *RPGRIP1L*^-/-^HEK293 cells (Fig. EV1B). To exclude *in vitro* artefacts, we also tested the role of RPGRIP1L in PCD regulation *in vivo*. In *Rpgrip1l*^-/-^ murine limb buds at embryonic day (E)12.5, PCD was significantly enhanced (Fig. 2B). In summary, our *in vitro* and *in vivo* data reveal that RPGRIP1L regulates PCD in vertebrates.

**Fig. 2:**
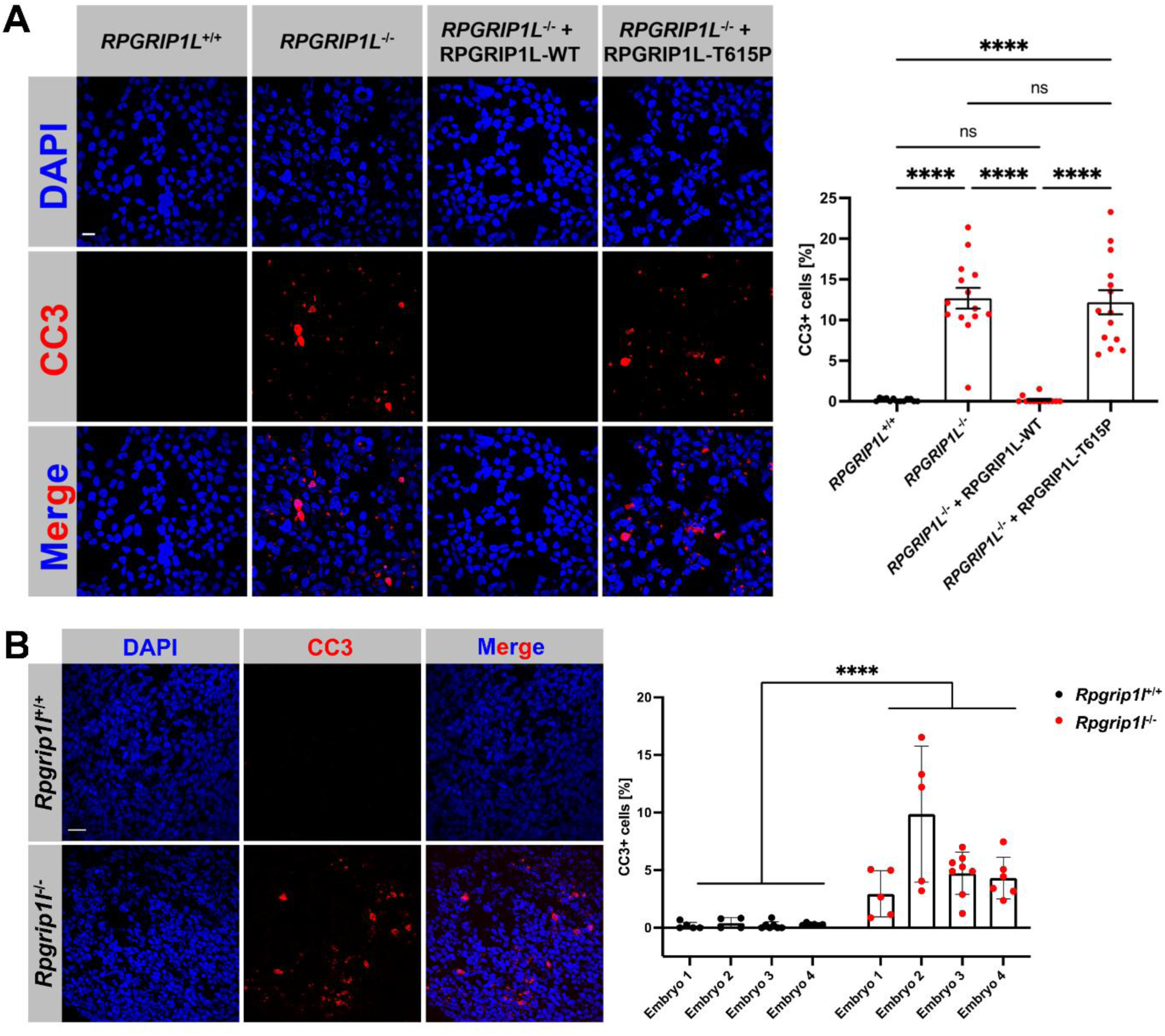
RPGRIP1L deficiency leads to an enhanced PCD in human cells and in mouse limb buds. (**A**) Immunofluorescence on serum-deprived HEK293 cells. Cell nuclei are stained in blue by DAPI. Cells undergoing PCD are marked in red by CC3. The scale bar (in white) represents a length of 20 μm. At least 1600 cells per genotype and transfection condition were used for these measurements, pooled from three replicates. Data shown as mean ± s.e.m. Statistical evaluation was performed by using Brown-Forsythe and Welch ANOVA test with Dunnett’s T3 multiple comparisons test (*RPGRIP1L*^+/+^ vs. *RPGRIP1L*^-/-^: **** *P* <0.0001; *RPGRIP1L*^+/+^ vs. *RPGRIP1L*^-/-^ + RPGRIP1L-T615P: **** *P* <0.0001; *RPGRIP1L*^- /-^ vs. *RPGRIP1L*^-/-^ + RPGRIP1L-WT: **** *P* <0.0001; *RPGRIP1L*^-/-^ + RPGRIP1L- WT vs. *RPGRIP1L*^-/-^ + RPGRIP1L-T615P: **** *P* <0.0001). ns = not significant. (**B**) Immunofluorescence on mouse limb buds which were obtained from *Rpgrip1l*^+/+^ (*n* = 4) and *Rpgrip1l*^-/-^ (*n* = 4) embryos at E12.5. Cell nuclei are stained in blue by DAPI. Cells undergoing PCD are marked in red by CC3. The scale bar (in white) represents a length of 30 μm. At least 1700 cells per limb bud were used for quantification, pooled from three replicates. Data shown as mean ± s.e.m. Statistical evaluation was performed by using two-tailed Mann-Whitney U test. Asterisks denote statistical significance (*** *P* = 0.0001).

### RPGRIP1L controls PCD by regulating proteasomal activity at the base of primary cilia

Our former studies demonstrated that RPGRIP1L deficiency results in impaired ciliary gating, in decreased autophagy and in reduced proteasomal activity at the ciliary base (ciliary proteasome) ^20–23^. These defects can be rescued by the use of the flavonoid eupatilin (EUP), the autophagy activator rapamycin (RAP) and the proteasome inducer sulforaphane (SFN), respectively ^21–23^. To evaluate whether these known functions of RPGRIP1L are associated with its PCD-regulating function, we treated *Rpgrip1l*^-/-^ NIH3T3 cells with EUP, RAP and SFN and quantified the percentage of cells undergoing PCD. The treatment with EUP or with RAP did not improve PCD in serum-deprived *Rpgrip1l*^-/-^ NIH3T3 cells, whereas the application of SFN completely rescued PCD in these cells (Fig. 3A) suggesting that RPGRIP1L governs PCD by its proteasome-regulating function. If this hypothesis is true, the inhibition of proteasomal activity at the ciliary base should cause an increase in PCD. As the regulation of the ciliary proteasome remains largely elusive and as there is no ciliary proteasome-specific inhibitor yet, we used the well-known proteasome inhibitor MG132 which concomitantly inhibits the activity of all proteasomes within a cell. The treatment of serum-deprived wild-type NIH3T3 cells with MG132 resulted in an increased PCD (Fig. 3B) supporting our hypothesis. To test whether MG132 treatment affects PCD due to the inhibition of the ciliary proteasome or due to the inhibition of the overall cellular proteasomal activity, we treated serum-deprived *Rpgrip1l*^-/-^NIH3T3 cells with MG132. We found that the percentage of CC3-positive *Rpgrip1l*^-/-^ NIH3T3 cells was not increased by the treatment with MG132 (Fig. 3C) demonstrating that RPGRIP1L controls PCD by regulating proteasomal activity at the ciliary base. By using the Proteasome Sensor Vector, we previously showed that RPGRIP1L regulates proteasomal activity exclusively at the ciliary base in mouse embryonic fibroblasts (MEFs) ^23^. Here, we performed this experiment in HEK293 cells and came to the same conclusion (Fig. 4A). The overall cellular proteasomal activity was unaltered in *RPGRIP1L*^-/-^ HEK293 cells (Fig. 4B). To confirm the regulation of PCD by the proteasome, serum-deprived wild-type HEK293 cells were treated with MG132. In line with the results obtained from NIH3T3 cells, MG132-treated HEK293 cells exhibit enhanced PCD (Fig. 4C). Furthermore, a treatment of serum-deprived *RPGRIP1L*-negative HEK293 cells with SFN rescued PCD (Fig. 4C).

**Fig. 3:**
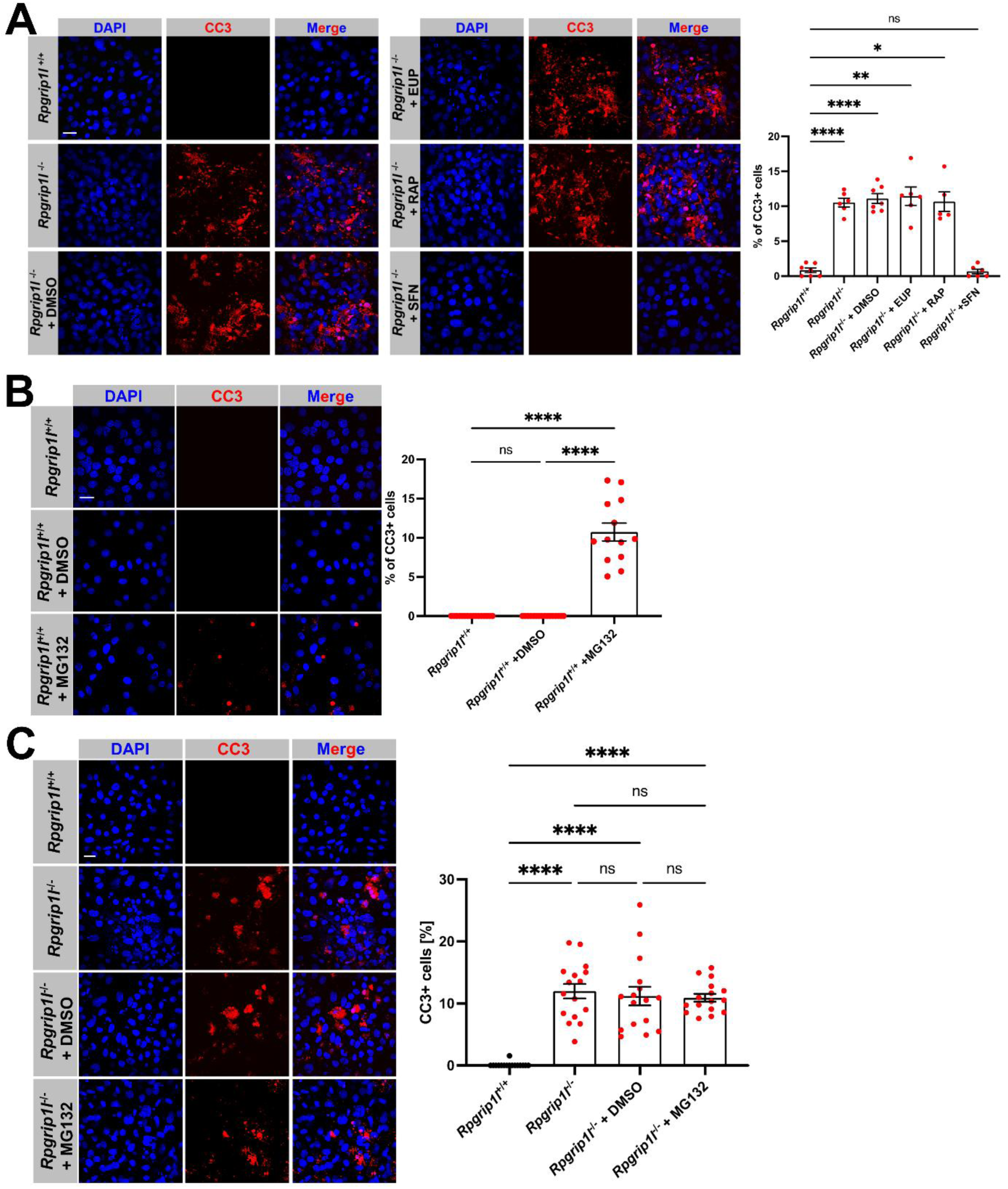
RPGRIP1L controls PCD by regulating the activity of the ciliary proteasome in NIH3T3 cells. (**A-C**) Immunofluorescence on serum-starved NIH3T3 cells. Cell nuclei are stained in blue by DAPI. Cells undergoing PCD are marked in red by CC3. The scale bars (in white) represent a length of 20 μm. (**A**) At least 580 cells per genotype and treatment were used for these measurements, pooled from three replicates. Data shown as mean ± s.e.m. Statistical evaluation was performed by using Brown-Forsythe and Welch ANOVA test with Dunnett’s T3 multiple comparisons test (*Rpgrip1l*^+/+^ vs. *Rpgrip1l*^-/-^: **** *P* <0.0001; *Rpgrip1l*^+/+^ vs. *Rpgrip1l*^-/-^ + DMSO: **** *P* <0.0001; *Rpgrip1l*^+/+^ vs. *Rpgrip1l*^-/-^ + EUP: ** *P* = 0.0025; *Rpgrip1l*^+/+^ vs. *Rpgrip1l*^-/-^ + RAP: ** *P* = 0.0186; *Rpgrip1l*^-/-^ vs. *Rpgrip1l*^-/-^ + SFN: **** *P* <0.0001; *Rpgrip1l*^-/-^ + DMSO vs. *Rpgrip1l*^-/-^ + SFN: **** *P* <0.0001; *Rpgrip1l*^-/-^ + EUP vs. *Rpgrip1l*^-/-^ + SFN: ** *P* = 0.0022; *Rpgrip1l*^-/-^ + RAP vs. *Rpgrip1l*^-/-^ + SFN: * *P* = 0.0174). ns = not significant. (**B**) At least 780 cells per treatment were used for quantification, pooled from three replicates. Data shown as mean ± s.e.m. Statistical evaluation was performed by using ordinary one-way ANOVA with Tukey’s multiple comparison test. Asterisks denote statistical significance (*Rpgrip1l*^+/+^ vs. *Rpgrip1l*^+/+^ + MG132: **** *P* <0.0001; *Rpgrip1l*^+/+^ + DMSO vs. *Rpgrip1l*^+/+^ + MG132: **** *P* <0.0001). ns = not significant. (**C**) At least 940 cells per genotype and treatment were used for quantification, pooled from three replicates. Data shown as mean ± s.e.m. Statistical evaluation was performed by using Brown-Forsythe and Welch ANOVA test with Dunnett’s T3 multiple comparisons test. Asterisks denote statistical significance (*Rpgrip1l*^+/+^ vs. *Rpgrip1l*^-/-^: **** *P* <0.0001; *Rpgrip1l*^+/+^ vs. *Rpgrip1l*^-/-^ + DMSO: **** *P* <0.0001; *Rpgrip1l*^+/+^ vs. *Rpgrip1l*^-/-^ + MG132: **** *P* <0.0001). ns = not significant.

**Fig. 4:**
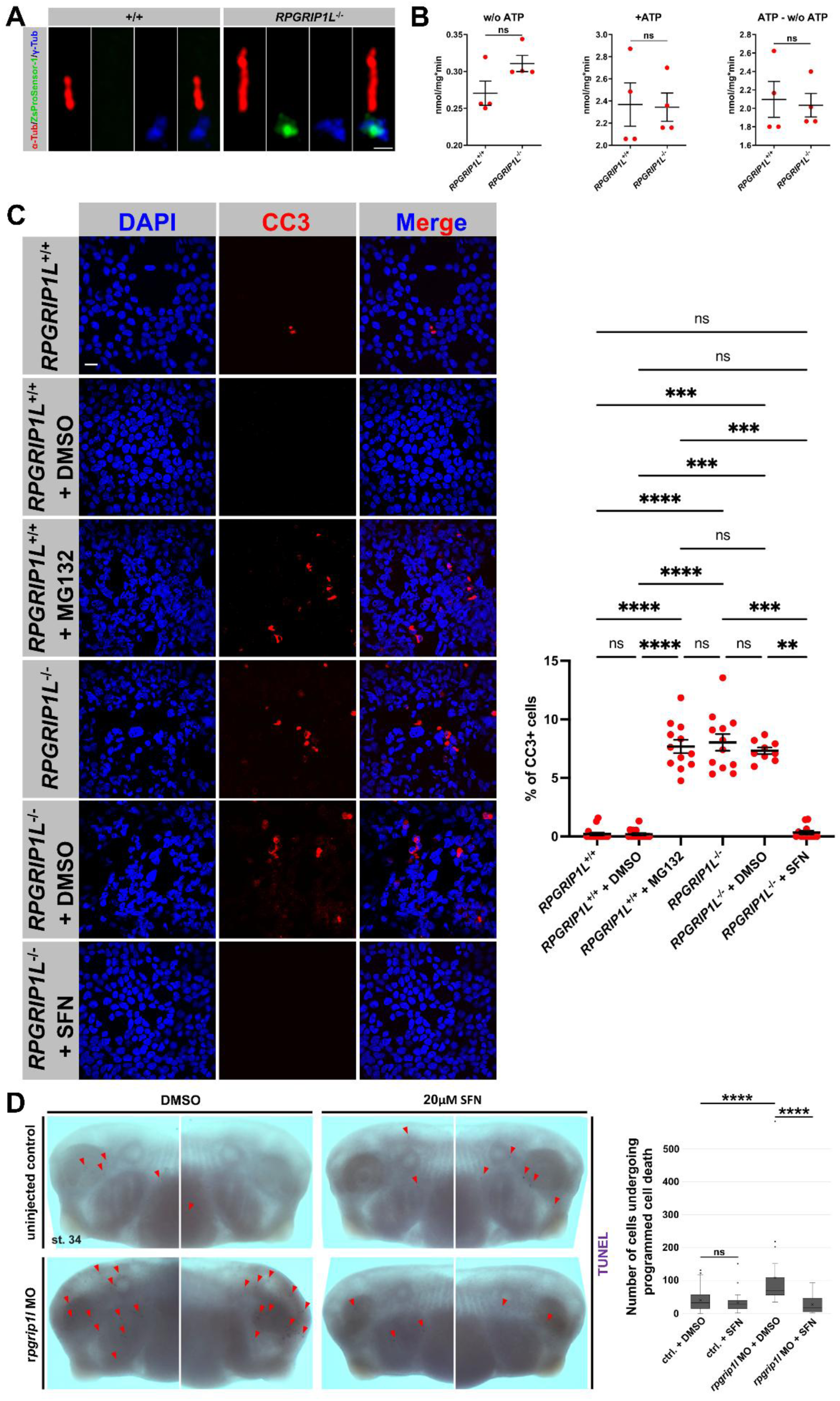
Restoration of proteasomal activity under RPGRIP1L deficiency rescues PCD *in vitro* and *in vivo*. (**A**) Fluorescence-based proteasome activity assay on serum-starved HEK293 cells. The ciliary axoneme is stained in green by acetylated α-tubulin and the BB in blue by γ-tubulin. The ZsProSensor-1 protein signal shines green. The scale bar (in white) represents a length of 1 μm. (**B**) Proteasomal activity was quantified in total cell lysates of serum-starved HEK293 cells. The activity of the 20S proteasomal subunit was measured without ATP. The activity of the 20S proteasomal subunit and the 26S proteasome together was quantified after ATP addition. The activity of the 26S proteasome was determined by subtracting the value measured for the activity of the 20S proteasomal subunit from the value measured for the activity of the 20S proteasomal subunit and the 26S proteasome together. ns = not significant. (**C**) Immunofluorescence on serum-starved HEK293 cells. Cell nuclei are stained in blue by DAPI. Cells undergoing PCD are marked in red by CC3. The scale bars (in white) represent a length of 20 μm. At least 1000 cells per genotype and treatment were used for these measurements, pooled from three replicates. Data shown as mean ± s.e.m. Statistical evaluation was performed by using Kruskal-Wallis test with Dunn’s multiple comparisons test (*RPGRIP1L*^+/+^ vs. *RPGRIP1L*^+/+^ + MG132: **** *P* <0.0001; *RPGRIP1L*^+/+^ vs. *RPGRIP1L* ^-/-^: **** *P* <0.0001; *RPGRIP1L*^+/+^ vs. *RPGRIP1L*^-/-^ + DMSO: *** *P* = 0.0002; *RPGRIP1L*^+/+^ + DMSO vs. *RPGRIP1L*^+/+^ + MG132: **** *P* <0.0001; *RPGRIP1L*^+/+^ + DMSO vs. *RPGRIP1L*^- /-^: **** *P* <0.0001; *RPGRIP1L*^+/+^ + DMSO vs. *RPGRIP1L*^-/-^ + DMSO: *** *P* = 0.0004; *RPGRIP1L*^+/+^ + MG132 vs. *RPGRIP1L*^-/-^ + SFN: *** *P* = 0.0005; *RPGRIP1L*^-/-^ vs. *RPGRIP1L*^- /-^ + SFN: *** *P* = 0.0003; *RPGRIP1L*^-/-^ + DMSO vs. *RPGRIP1L*^-/-^ + SFN: ** *P* = 0.0026). ns = not significant. (**D**) TUNEL assay on control + DMSO (*n* = 35), control + SFN (*n* = 39), *rpgrip1l* MO + DMSO (*n* = 23) and *rpgrip1l* MO + SFN (*n* = 33) *Xenopus* embryos at stage 34, pooled from three replicates. Data shown as mean ± s.e.m. Statistical evaluation was performed by using two-tailed Mann-Whitney U test. Asterisks denote statistical significance (control + DMSO vs. *rpgrip1l* MO + DMSO: **** *P* <0.0001; *rpgrip1l* MO + DMSO vs. *rpgrip1l* MO + SFN: **** *P* <0.0001). ns = not significant.

Next, we tested whether SFN could also rescue PCD *in vivo*. As the treatment of mouse embryos *in utero* is difficult and time-consuming, we used an alternative vertebrate model system – the clawed frog *Xenopus laevis*. We generated *rpgrip1l* knockdown *Xenopus* embryos by using morpholino oligonucleotides (MOs) and investigated cell death by performing TdT-mediated dUTP nick end labelling (TUNEL) staining. In *rpgrip1l*-deficient *Xenopus* embryos, PCD was significantly increased at stage 34 (Fig. 4D). SFN was administered to *rpgrip1l*-deficient *Xenopus* embryos and rescued PCD (Fig. 4D) confirming the results obtained in the *in vitro* rescue experiments.

### RPGRIP1L controls PCD by regulating the proteasomal degradation of Modulator of Apoptosis 1 (MOAP1) at the ciliary base

As mentioned above, the percentage of ciliated cells was decreased in *Rpgrip1l*^-/-^ NIH3T3 cells (Fig. 1C). In search for the mechanism underlying the regulation of PCD by RPGRIP1L, we tested whether the treatment with SFN rescues PCD in *Rpgrip1l*^-/-^ NIH3T3 cells by increasing the percentage of ciliated cells. As SFN did not increase the percentage of ciliated cells (Fig. EV2), we rejected this hypothesis. Since RPGRIP1L is a key regulator of TZ assembly and hence controls ciliary gating and ciliary protein composition, we hypothesised that the loss of RPGRIP1L affects the localisation of cell death receptors in the ciliary membrane. According to our knowledge, it is unknown whether the main death receptors, such as Tumor Necrosis Factor Receptor 1 (TNF-R1) or Tumor Necrosis Factor Related Apoptosis-Inducing Ligand Receptor 1 (TRAIL-R1), show a ciliary localisation. So, we checked for ciliary presence of these receptors by using immunofluorescence. In serum-deprived wild-type NIH3T3 cells, we detected TNF-R1 at the basal body and at daughter centriole but not within cilia (Fig. EV3A). In contrast, TRAIL-R1 was located at the basal body and daughter centriole and, in addition, along the whole axoneme (Fig. EV3B). We quantified the amount of TNF-R1 at the basal body as well as at daughter centriole and of TRAIL-R1 at the basal body, at daughter centriole and along the entire axoneme and could not find any difference between wild-type and *Rpgrip1l*^-/-^NIH3T3 cells (Fig. EV3A,B). This precluded that the increased PCD observed in the absence of RPGRIP1L is caused by alterations at the level of the cell death receptors.

To find potential interaction partners of RPGRIP1L that are associated with PCD, we performed a yeast two-hybrid screen. Among the twelve identified potential interaction partners of RPGRIP1L (Fig. EV4A), we assessed MOAP1 (alias MAP1 or PNMA4) as the only candidate which might be involved in the RPGRIP1L-controlled regulation of PCD. We based our theory on two points: 1) MOAP1 induces apoptosis by interacting with the BCL2 Associated X (BAX) protein, an important regulator of apoptosis ^24^. 2) It was reported that the proteasomal degradation of MOAP1 is deeply involved in the regulation of apoptosis ^25^. To test our theory, we performed a series of experiments. First of all, we confirmed the interaction of RPGRIP1L with MOAP1 by using a co-immunoprecipitation assay. As murine full-length RPGRIP1L could not be stably expressed, we used a FLAG-tagged RPGR-interacting domain (RID) of RPGRIP1L and a MYC-tagged full-length MOAP1 for transient overexpression. By co-immunoprecipitating these fusion proteins, we confirmed the interaction of RPGRIP1L with MOAP1 (Fig. 5A). To unravel in which subcellular region these proteins interact with each other, we performed an *in situ* proximity ligation assay (*in situ* PLA) as previously described ^26^. By using this assay, a fluorescence signal (PLA signal) reveals the intracellular interaction site. In all cases, we detected PLA signals exclusively at the ciliary base of wild-type NIH3T3 cells (13 out of 13 ciliated cells) but not in *Rpgrip1l*^-/-^ NIH3T3 cells (0 out of 12 ciliated cells) (Fig. 5B). To elucidate the subciliary localisation of MOAP1, we performed super-resolution microscopy. By using Airyscan super-resolution microscopy (image scanning microscopy) ^27,28^, we showed that MOAP1 localises to the basal body (Fig. 5C; Movie EV1). Moreover, we found MOAP1 at the daughter centriole. To achieve an even higher resolution and hence to evaluate the ciliary localisation of MOAP1 in more detail, we performed single-molecule localisation microscopy by using dSTORM ^29,30^. dSTORM revealed that MOAP1 protrudes into the basal body up to the transition zone elucidating the close proximity to RPGRIP1L (Fig. 5D; Movies EV2, EV3 and EV4). Since a higher amount of MOAP1 results in an increase of apoptosis ^24^, we quantified the amount of MOAP1 in *Rpgrip1l*^-/-^ NIH3T3 cells. We found that the overall cellular amount of MOAP1 was unaltered in *Rpgrip1l*^-/-^ NIH3T3 cells (Fig. EV4B). Furthermore, the amount of MOAP1 at the daughter centriole in *Rpgrip1l*^-/-^ NIH3T3 cells was not changed (Fig. EV4C). In contrast, we detected an enhanced amount of MOAP1 at the ciliary base in *Rpgrip1l*^-/-^ NIH3T3 cells (Fig. 5E). Importantly, this increased ciliary amount of MOAP1 was rescued by treating *Rpgrip1l*^-/-^ NIH3T3 cells with SFN (Fig. 5E). To test whether the enhanced ciliary amount of MOAP1 could be the reason for the significantly higher number of cells undergoing PCD in the absence of RPGRIP1L, we reduced the amount of MOAP1 by using RNA interference (RNAi). After the transfection of *Rpgrip1l*^-/-^ NIH3T3 cells with small interfering RNA (siRNA) against *Moap1*, we quantified a strongly reduced amount of MOAP1 at the ciliary base (Fig. 5F) and a decreased percentage of cells undergoing PCD (Fig. 5G). These findings demonstrate that the regulation of PCD by RPGRIP1L is realised via the proteasomal degradation of MOAP1.

**Fig. 5:**
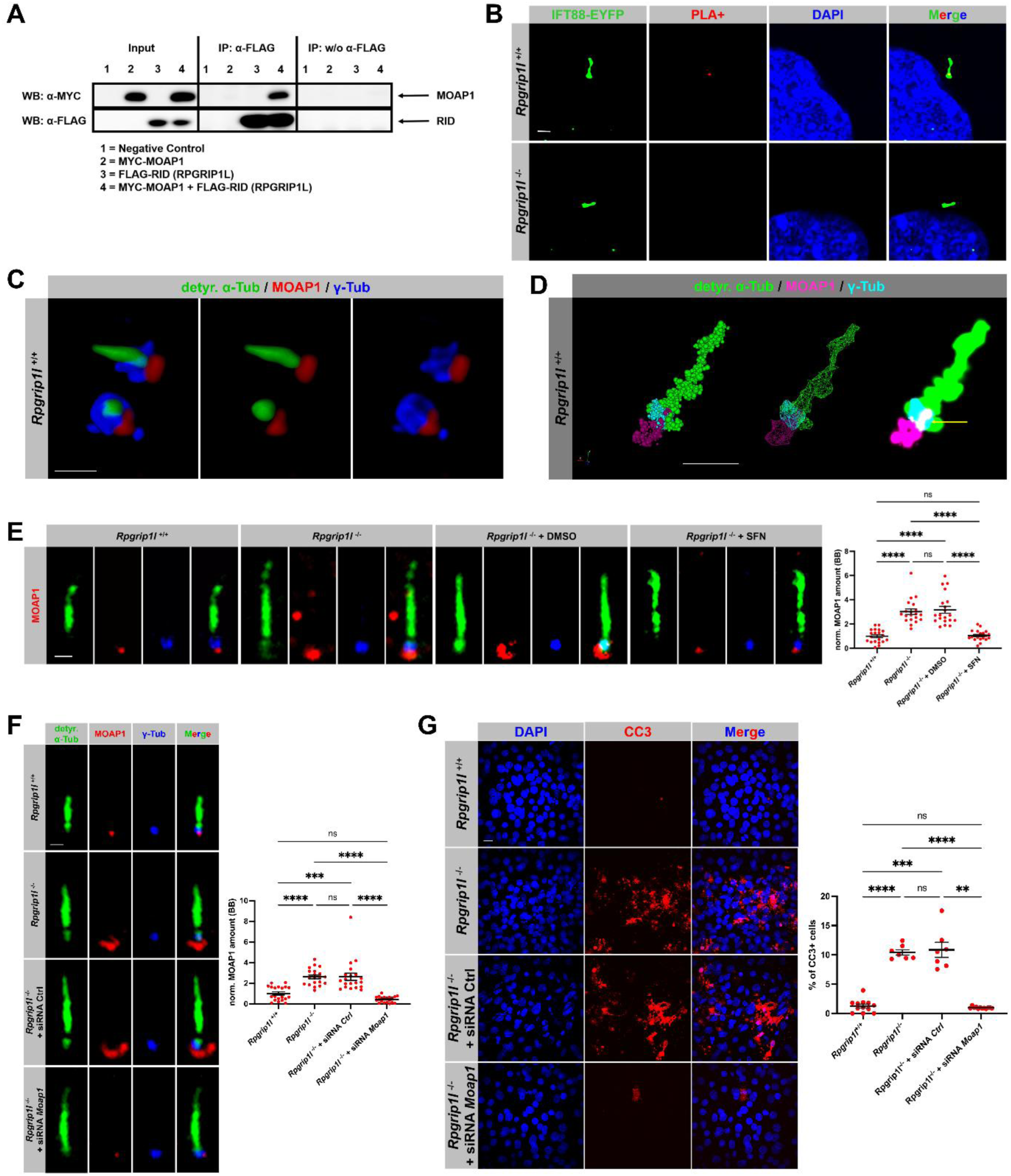
RPGRIP1L governs PCD via ensuring the proteasomal degradation of MOAP1 at the ciliary base. (**A**) Coimmunoprecipitation in NIH3T3 cells. MYC-tagged MOAP1 full-length protein and FLAG-tagged RID (domain of RPGRIP1L) were transiently overexpressed and tested for interaction by coimmunoprecipitation from total cell lysates. Immunoprecipitation assays were performed by using an anti-FLAG antibody. IP, immunoprecipitation; WB, western blot. (**B**) *In situ* proximity ligation assay (*in situ* PLA) on serum-starved NIH3T3 cells. Cell nuclei are stained in blue by DAPI and cilia are marked by the product of a transiently transfected IFT88-EYFP construct. The PLA+ signal is shown in red. The scale bar (in white) represents a length of 2 μm. (**C**) Immunofluorescence on serum-starved NIH3T3 cells. Super-resolution image was acquired by using Airyscan microscopy. The ciliary axoneme is labelled in green by detyrosinated α-tubulin, the basal body is marked in blue by γ-tubulin. MOAP1 is shown in red. The scale bar (in white) represents a length of 5 μm. (**D**) Immunofluorescence on serum-starved NIH3T3 cells. Super-resolution image was acquired by using dSTORM. The ciliary axoneme is labelled in green by detyrosinated α-tubulin, the basal body is marked in turquoise by γ-tubulin. MOAP1 is shown in magenta. The scale bar (in white) represents a length of 1 μm. The depicted cilium is shown in three different modes of representation – the point cloud mode (on the left), the cluster mode (in the middle) and point splatting mode (on the right). In the point splatting mode, the yellow arrow points to the portion of MOAP1 that protrudes into the basal body up to the transition zone (shown in white). (**E**) Immunofluorescence on serum-starved NIH3T3 cells. The ciliary axoneme is stained in green by detyrosinated α-tubulin and the BB in blue by γ-tubulin. The scale bar (in white) represents a length of 1 μm. 20 cilia per genotype were used for quantification, pooled from three replicates. Data shown as mean ± s.e.m. Statistical evaluation was performed by using Kruskal-Wallis test with Dunn’s multiple comparisons test. Asterisks denote statistical significance (*Rpgrip1l*^+/+^ vs. *Rpgrip1l*^-/-^: **** *P* <0.0001; *Rpgrip1l*^+/+^ vs. *Rpgrip1l*^-/-^ + DMSO: **** *P* <0.0001; *Rpgrip1l*^-/-^ vs. *Rpgrip1l*^-/-^ + SFN: **** *P* <0.0001; *Rpgrip1l*^-/-^ + DMSO vs. *Rpgrip1l*^-/-^ + SFN: **** *P* <0.0001). ns = not significant. (**F**) Immunofluorescence on serum-starved NIH3T3 cells. The ciliary axoneme is stained in green by detyrosinated α-tubulin and the BB in blue by γ-tubulin. The scale bar (in white) represents a length of 1 μm. 20 cilia per genotype were used for quantification, pooled from three replicates. Data shown as mean ± s.e.m. Statistical evaluation was performed by using Kruskal-Wallis test with Dunn’s multiple comparisons test. Asterisks denote statistical significance (*Rpgrip1l*^+/+^ vs. *Rpgrip1l*^-/-^: **** *P* <0.0001; *Rpgrip1l*^+/+^ vs. *Rpgrip1l*^-/-^ + siRNA Ctrl.: **** *P* = 0.0009; *Rpgrip1l*^-/-^ vs. *Rpgrip1l*^-/-^ + siRNA *Moap1*: **** *P* <0.0001; *Rpgrip1l*^- /-^ siRNA Ctrl. vs. *Rpgrip1l*^-/-^ + siRNA *Moap1*: **** *P* <0.0001). ns = not significant; Ctrl., control. (**G**) Immunofluorescence on serum-starved NIH3T3 cells. Cell nuclei are stained in blue by DAPI. Cells undergoing PCD are marked in red by CC3. The scale bar (in white) represents a length of 10 μm. At least 800 cells per genotype and treatment were used for quantification, pooled from three replicates. Data shown as mean ± s.e.m. Statistical evaluation was performed by using Brown-Forsythe and Welch ANOVA test with Dunnett’s T3 multiple comparisons test. Asterisks denote statistical significance (*Rpgrip1l*^+/+^ vs. *Rpgrip1l*^-/-^: **** *P* <0.0001; *Rpgrip1l*^+/+^ vs. *Rpgrip1l*^-/-^ + siRNA Ctrl.: *** *P* = 0.0009; *Rpgrip1l*^-/-^ vs. *Rpgrip1l*^- /-^ + siRNA *Moap1*: **** *P* <0.0001; *Rpgrip1l*^-/-^ + siRNA Ctrl. vs. *Rpgrip1l*^-/-^ + siRNA *Moap1*: ** *P* = 0.0013). ns = not significant; Ctrl., control.

### Beyond RPGRIP1L, CEP290 and TCTN1 govern PCD by controlling the activity of the ciliary proteasome

As at least 28 proteins are located at the TZ ^31^ and as RPGRIP1L acts as a central factor in vertebrate TZ assembly ^20^, the question arises whether other TZ proteins also exert an PCD-regulating function by controlling proteasomal activity at the ciliary base. To select promising candidates, we looked for TZ proteins for which it was previously shown that they regulate GLI3 processing. Since GLI3 is proteolytically processed into its repressor form by the ciliary proteasome ^23,32^, these proteins might regulate the proteasomal activity at the ciliary base. In mice, mutations in *Tctn1* or in *Cep290* result in an impaired GLI3 processing and hence in a reduced amount of the GLI3 repressor form ^33,34^. The same processing defect was detected in ciliopathy patient-derived cells with mutations in *CEP290* ^35^. Accordingly, we decided to investigate proteasomal activity in the absence of TCTN1 or CEP290 and used *Tctn1*^-/-^ and *Cep290*^-/-^ NIH3T3 cells we generated before ^20^. While the overall cellular proteasomal activity was unaltered in both (Fig. 6A,B), we found a reduced activity of the proteasome at the ciliary base in serum-deprived *Tctn1*^-/-^ NIH3T3 cells and in serum-deprived *Cep290*^-/-^ NIH3T3 cells (Fig. 6C). Next, we analysed if PCD is altered in the absence of TCTN1 or CEP290. Significantly more serum-deprived *Tctn1*^-/-^ NIH3T3 cells as well as *Cep290*^-/-^ NIH3T3 cells were positive for CC3 than wild-type NIH3T3 cells (Fig. 7A,B). Importantly, the treatment of *Tctn1*^-/-^ NIH3T3 cells or *Cep290*^-/-^ NIH3T3 cells with SFN rescued PCD (Fig. 7A,B) demonstrating that TCTN1 and CEP290 control PCD by regulating proteasomal activity at the ciliary base. In contrast to RPGRIP1L, TCTN1 and CEP290, it was not reported so far that Inversin (INVS), a protein that localises to the ciliary TZ of NIH3T3 cells ^20^, is involved in GLI3 processing. In line with that, our data reveal that neither proteasomal activity nor PCD was altered in *Invs*^-/-^ NIH3T3 cells (Fig. EV5A-C). To test whether SFN also rescues PCD under CEP290 deficiency in our *in vivo* model, we knocked down *cep290* in *Xenopus* embryos by MO injection and analysed PCD at stage 34. The generated *cep290* morphants displayed increased PCD (Fig. 7C). PCD was rescued in *cep290* morphant *Xenopus laevis* embryos by the treatment with SFN (Fig. 7C). These data demonstrate that the regulation of PCD by governing the activity of the ciliary proteasome is not RPGRIP1L-specific and indicate that several TZ proteins might be able to control PCD in the same way.

**Fig. 6:**
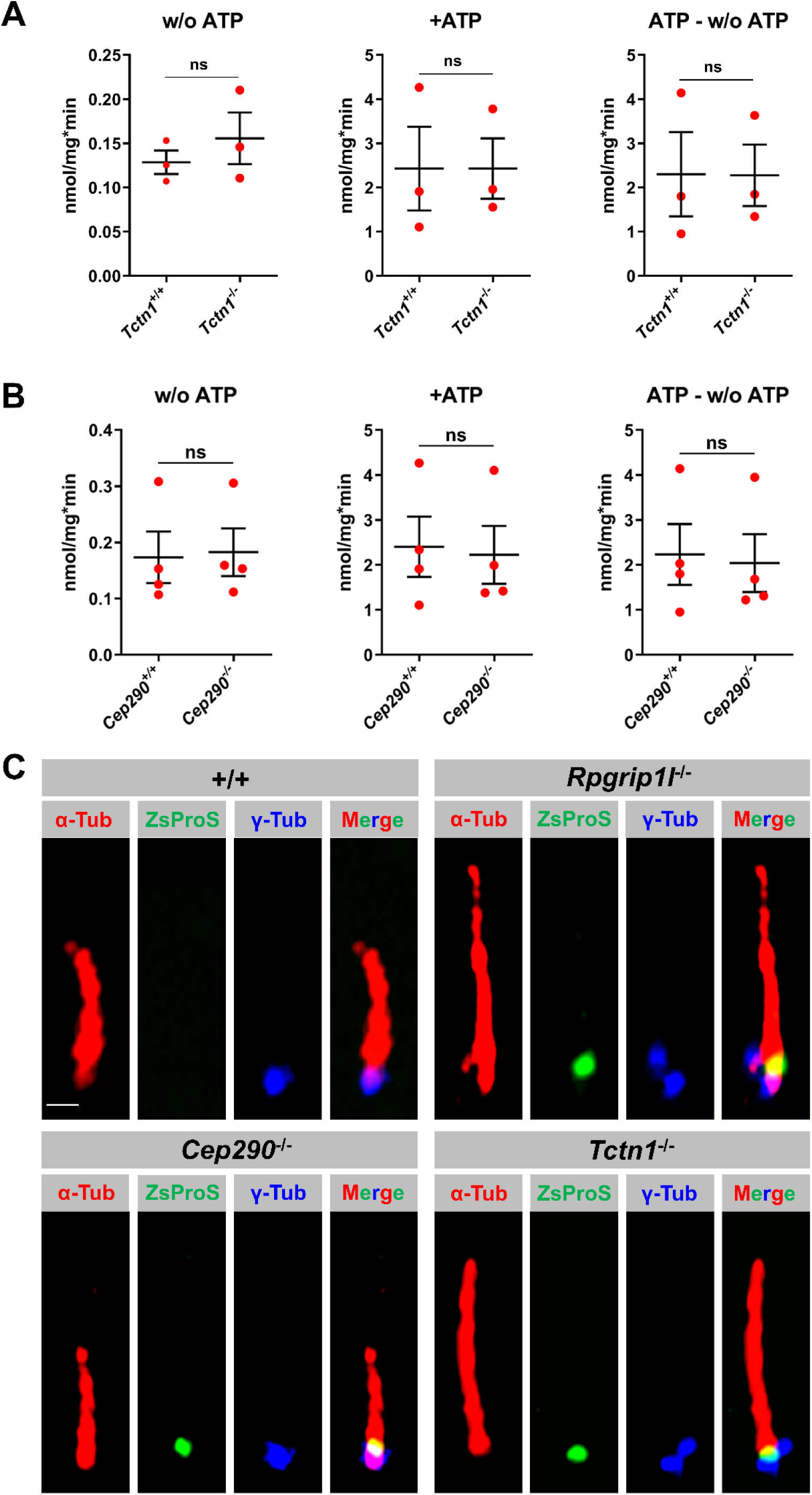
TCTN1 and CEP290 regulate the activity of the ciliary proteasome. (**A, B**) Proteasomal activity was quantified in total cell lysates of serum-starved NIH3T3 cells. The activity of the 20S proteasomal subunit was measured without ATP. The activity of the 20S proteasomal subunit and the 26S proteasome together was quantified after ATP addition. The activity of the 26S proteasome was determined by subtracting the value measured for the activity of the 20S proteasomal subunit from the value measured for the activity of the 20S proteasomal subunit and the 26S proteasome together. ns = not significant. (**C**) Fluorescence-based proteasome activity assay on serum-starved NIH3T3 cells. The ciliary axoneme is stained in green by acetylated α-tubulin and the BB in blue by γ-tubulin. The ZsProSensor-1 protein signal shines green. The scale bar (in white) represents a length of 0.5 μm.

**Fig. 7:**
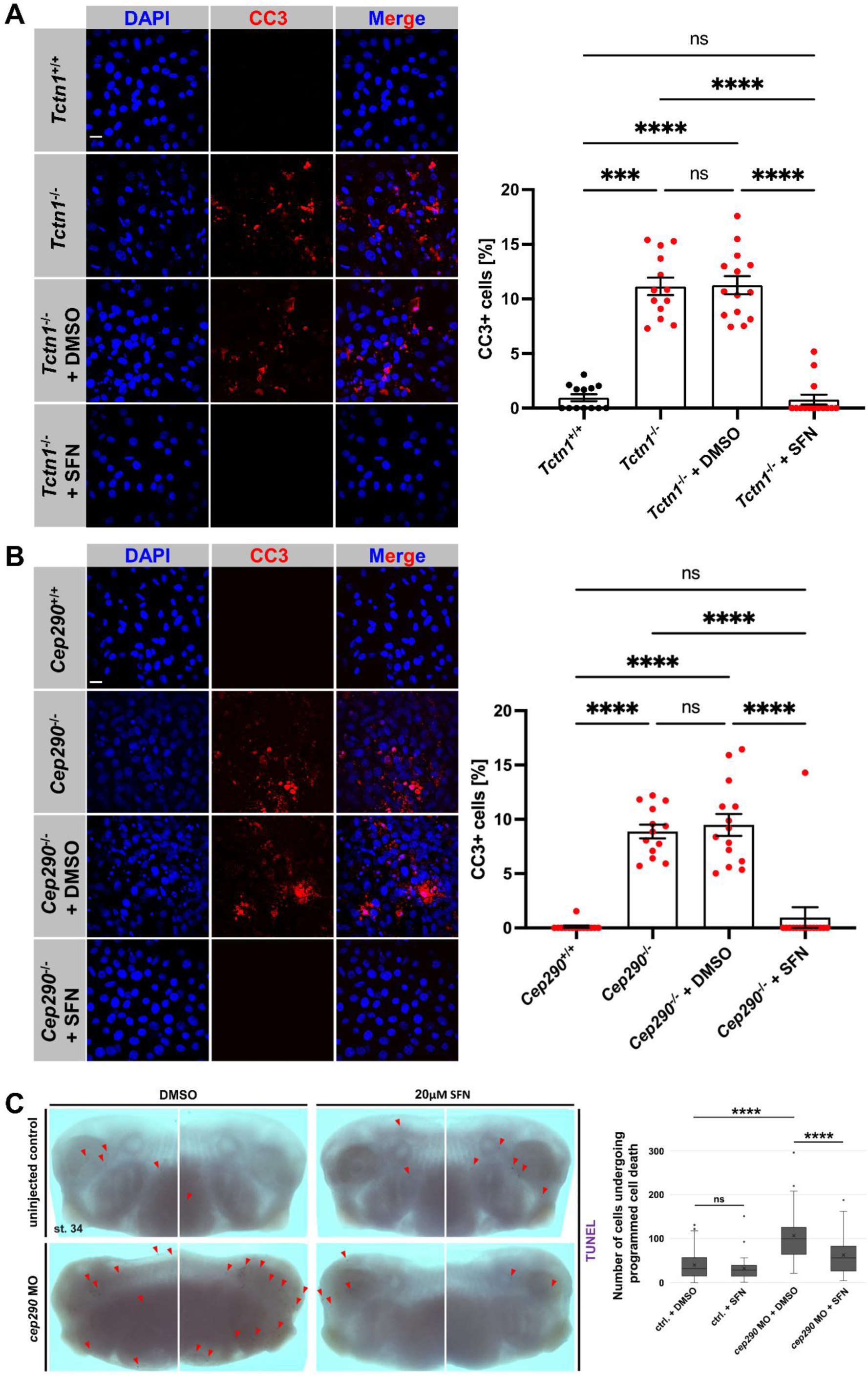
TCTN1 and CEP290 control PCD by regulating proteasomal activity. (**A, B**) Immunofluorescence on serum-starved NIH3T3 cells. Cell nuclei are stained in blue by DAPI. Cells undergoing PCD are marked in red by CC3. The scale bars (in white) represent a length of 20 μm. (**A**) At least 650 cells per genotype and treatment were used for these measurements, pooled from three replicates. Data shown as mean ± s.e.m. Statistical evaluation was performed by using Kruskal-Wallis test with Dunn’s multiple comparisons test. Asterisks denote statistical significance (*Tctn*^+/+^ vs. *Tctn1*^-/-^: **** *P* = 0.0001; *Tctn1*^+/+^ vs. *Tctn1*^-/-^ + DMSO: **** *P* <0.0001; *Tctn1*^-/-^ vs. *Tctn1*^-/-^ + SFN: **** *P* <0.0001; *Tctn1*^-/-^ + DMSO vs. *Tctn1*^-/-^ + SFN: **** *P* <0.0001). ns = not significant. (**B**) At least 790 cells per genotype and treatment were used for these measurements, pooled from three replicates. Data shown as mean ± s.e.m. Statistical evaluation was performed by using Kruskal-Wallis test with Dunn’s multiple comparisons test. Asterisks denote statistical significance (*Cep290*^+/+^ vs. *Cep290* ^-/-^: **** *P* <0.0001; *Cep290*^+/+^ vs. *Cep290*^-/-^ + DMSO: **** *P* <0.0001; *Cep290*^-/-^ vs. *Cep290*^-/-^ + SFN: **** *P* <0.0001; *Cep290*^-/-^ + DMSO vs. *Cep290*^-/-^ + SFN: **** *P* <0.0001). ns = not significant. (**C**) TUNEL assay on control + DMSO (*n* = 35), control + SFN (*n* = 39), *cep290* MO + DMSO (*n* = 25) and *cep290* MO + SFN (*n* = 31) *Xenopus* embryos at stage 34, pooled from three replicates. Data shown as mean ± s.e.m. Statistical evaluation was performed by using two-tailed Mann-Whitney U test. Asterisks denote statistical significance (control + DMSO vs. *cep290* MO + DMSO: **** *P* <0.0001; *cep290* MO + DMSO vs. *cep290* MO + SFN: **** *P* <0.0001). ns = not significant.

## Discussion

Primary cilia are deeply involved in the regulation of several cellular processes, including proliferation, migration, differentiation etc., as they transduce signals from the environment into the cell’s interior. Thus, ciliary dysfunction results in the development of ciliopathies. Remarkably, several symptoms of ciliopathy syndromes come along with increased PCD ^36–38^. However, the cilia-mediated regulation of PCD remains only poorly understood ^9^. Our study contributes to a better understanding of this regulation. Apoptosis was originally believed to represent the only form of PCD. As a consequence of intensive research in the field of PCD, several different PCDs have been described in the last twenty-five years. Beyond apoptosis, three further types of PCDs exist: necroptosis, ferroptosis and pyroptosis (for details see ^39^). Recently, it was reported that primary cilia suppress Receptor Interacting Serine/Threonine Kinase 3 (RIPK3)-mediated necroptosis ^40^, whereas the role of primary cilia in the control of ferroptosis and pyroptosis remains elusive. According to the current state of knowledge, the mechanism described in this work refers to the control of apoptosis rather than of the other mentioned PCDs. We base this statement on the following facts: The number of CC3-positive and CC9-positive cells is increased in our cilia mutants (Fig. 1A,B). CC3 and CC9 are deeply involved in the apoptotic pathway. In its activated (cleaved) form, caspase-3 serves as an executioner protease functioning at the end of the apoptotic pathway. It is able to cleave a plethora of substrates, such as cell cycle regulators, structural proteins etc., and plays an essential role in the degradation of centrosomes. CC3 is activated, among other things, by CC9 ^39,41^. CC9 functions as part of the intrinsic (mitochondrial) apoptotic pathway but does not work in the extrinsic (death receptor-driven) apoptotic pathway. CC3 and CC9 are also involved in pyroptosis but not in necroptosis or ferroptosis ^39,42–44^. Furthermore, MOAP1, a central player in the mechanism described herein, has been so far associated with apoptosis but not with necroptosis, ferroptosis or pyroptosis ^45^. To further test whether this MOAP1-dependent mechanism by which primary cilia regulate PCD does not affect necroptosis, pyroptosis or ferroptosis, several marker proteins [e.g. RIPK3 (necroptosis), Mixed Lineage Kinase Domain Like Pseudokinase (MLKL; necroptosis), Gasdermin D (GSDMD; pyroptosis), Gasdermin D (GSDME; pyroptosis)] and inhibitors [e.g. ferrostatin-1 (ferroptosis), liproxstatin-1 (ferroptosis)] could be analysed in *Rpgrip1l*^-/-^ cells.

Previous studies showed that MOAP1 localises to the cytoplasm and is enriched at the outer mitochondrial membrane ensuring the mitochondrial accumulation of tBID, the activated form of BID that interacts with the key apoptosis regulators BAX and BAD ^24,46,47^. In this way, MOAP1 plays an important role in the intrinsic apoptotic pathway. Our data suggest a model in which MOAP1 localises to the base of primary cilia where it gets continuously degraded by the ciliary proteasome (Fig. 8). In the absence of certain transition zone proteins, such as RPGRIP1L or CEP290, the activity of the ciliary proteasome is impaired causing an accumulation of MOAP1 at the ciliary base. The increased ciliary amount of MOAP1 induces PCD (as discussed above, most likely apoptosis). Our findings raise the question of why the mitochondrial protein MOAP1 is recruited to the ciliary base. Interestingly, it was shown before that MOAP1 is not only involved in the intrinsic apoptotic pathway but also in the extrinsic, death receptor-dependent apoptotic pathway interacting with death receptors, such as TNF-R1 and TRAIL-R1 ^48^. Subsequently to the stimulation of these death receptors, MOAP1 is prevented from proteasomal degradation and hence able to activate BAX ^25^. The activation of BAX results in the release of cytochrome C from the mitochondria into the cytosol leading to the formation of the apoptosome which in turn activates caspase-9. In line with these previously published data, we quantified an increase in the percentage of CC9-positive NIH3T3 cells in the absence of RPGRIP1L (Fig. 1B). Importantly, our data show that cell death receptors localise at the ciliary base (TNF-R1 and TRAIL-R1) and along the whole cilium (TRAIL-R1) (Fig. EV3A,B). Based on these findings, we hypothesise that MOAP1 localises to primary cilia in order to interact with ciliary death receptors. Further, we suggest that if the proteasomal degradation of MOAP1 at the ciliary base is impeded, MOAP1 is transported from the ciliary base to the mitochondria. If there is no apoptotic stimulus, MOAP1 gets continually degraded by the ciliary proteasome and is not able to activate BAX in the mitochondria. Combining our novel findings with previously published data, we favour a scenario in which the ciliary localisation of MOAP1 is beneficial as MOAP1 localises close to the death receptors in the ciliary membrane and as MOAP1 can be degraded by the ciliary proteasome on the spot without any translocation. The mechanism suggested by our results represents another example for the vibrant crosstalk between primary cilia and mitochondria ^49–53^.

**Fig. 8:**
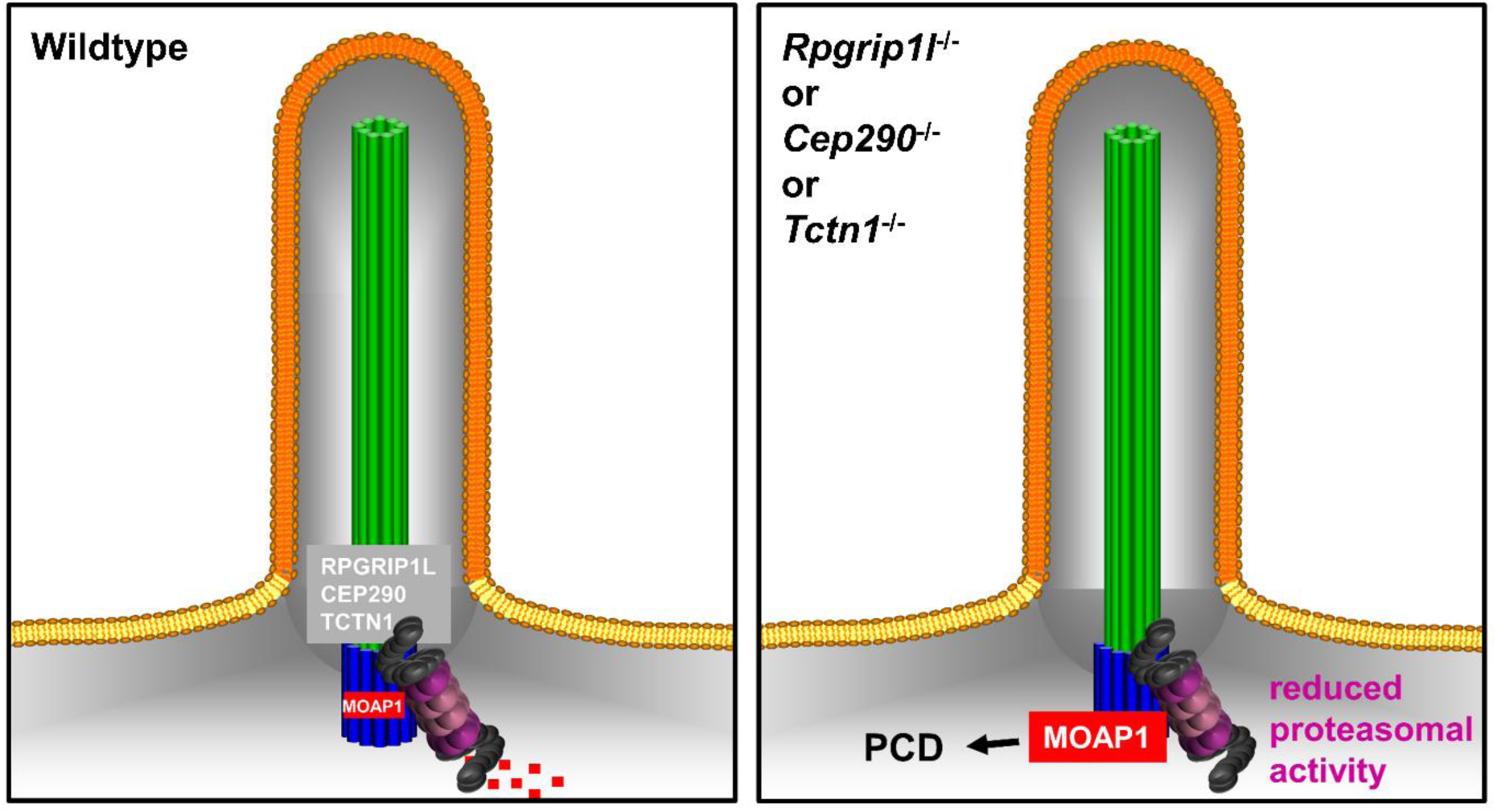
Graphical abstract of the mechanism by which primary cilia regulate PCD. In the wildtype, RPGRIP1L, CEP290 and TCTN1 positively regulate the activity of the ciliary proteasome. The ciliary proteasome constantly degrades MOAP1 at the ciliary base. In the absence of RPGRIP1L, CEP290 or TCTN1, the activity of the ciliary proteasome is severely reduced impeding the degradation of MOAP1 at the ciliary base. As a consequence, MOAP1 accumulates at the ciliary base and finally induces PCD.

In the absence of RPGRIP1L, the ciliary amount of MOAP1 was increased causing significantly enhanced PCD (Fig. 5D-F). In contrast to the amount of MOAP1 at the BB, the amount of MOAP1 at the daughter centriole and the overall cellular protein amount of MOAP1 were unaltered (Fig. EV3B,C) demonstrating that the ciliary proteasome serves as a decisive factor in the regulation of PCD. Accordingly, the percentage of ciliated cells is crucial for the induction of PCD in the absence of RPGRIP1L. A higher percentage of ciliated *Rpgrip1l*-negative NIH3T3 cells caused a stronger increase in PCD (Fig. 1C,D). In wild-type NIH3T3 cells, the degree of ciliation does not alter the percentage of cells undergoing PCD as the ciliary proteasome is working properly.

Beyond the control of PCD by RPGRIP1L via its regulation of the activity of the ciliary proteasome, additional mechanisms are conceivable by which RPGRIP1L governs PCD. We have previously shown that RPGRIP1L functions as a key factor in TZ assembly and ciliary gating ^20,21^. In the absence of RPGRIP1L, ciliary gating is impaired but can be rescued by the use of EUP ^21^. Ciliary gating governs ciliary protein composition, e.g. the amount of many receptors involved in various signalling pathways. For this reason, it is conceivable that more death receptors accumulate in the ciliary membrane when RPGRIP1L is missing. This could cause an increase of PCD by an overactivation of the extrinsic apoptotic pathway. However, our data contradict this possibility since EUP did not rescue PCD in *Rpgrip1l*^-/-^ NIH3T3 cells (Fig. 3A). In addition to TZ assembly and ciliary gating, RPGRIP1L regulates autophagy. Autophagy decreases by the loss of RPGRIP1L and is rescued by RAP in the absence of RPGRIP1L ^22^. It was reported before that reduced autophagy can result in increased PCD ^54^. However, treatment of *Rpgrip1l*^-/-^ NIH3T3 cells with the autophagy activator RAP did not rescue PCD (Fig. 3A) demonstrating that RPGRIP1L controls PCD independently of its autophagy-regulating function. Considering the different functions of RPGRIP1L, its proteasome-regulating function seems to be the only one involved in the regulation of PCD. Importantly, two more TZ proteins, TCTN1 and CEP290, also control PCD by regulating the activity of the ciliary proteasome (Fig. 8). These findings indicate a central role of the ciliary proteasome in the cilia-controlled regulation of PCD.

Apart from mutations in genes encoding TZ proteins, mutations in genes encoding proteins that are involved in the transport within cilia (intraflagellar transport; IFT) engenders a higher apoptosis rate in developing zebrafish eyes as well as in mouse brains and thyrocytes ^55–58^. Remarkably, some IFT mutants show an impaired GLI3 processing ^59–61^ arising the question whether an impaired IFT affects the activity of the ciliary proteasome. In a former study, we detected PSMA5, a member of the 20S proteasomal subunit, not only at the ciliary base but also along the whole axoneme ^23^ enhancing the likelihood that the IFT participates in the functional control of the ciliary proteasome.

Importantly, our data could provide insights into the cilia-mediated regulation of PCD in certain cancer cells. MOAP1 has previously shown to act as a tumour suppressor ^62^. A reduced amount of MOAP1 promotes tumorigenesis in several cancers. Notably, the inhibition of proteasomal activity with MG132 increases the amount of MOAP1 resulting in enhanced apoptosis in several cancer cell lines ^62^. In accordance with these findings, proteasomal activity is elevated in most cancer cell types ^3^. It is an exciting question for future studies whether the activity of the ciliary proteasome gets reduced when cells become malignant in order to prevent the cancer cells from undergoing apoptosis. Potentially, our results pave the way for the development of novel treatments not only to tackle ciliopathies but also cancer.

## Methods

### Animal husbandry and ethics statement

The origin of the mouse breeding was acquired from Charles River Laboratories, Sulzfeld, Germany. All mice (*Mus musculus*) used in this study were maintained in the C3H background and housed at 22-24°C on a 12/12 hours dark-light cycle with food and water *ad libitum*. All mouse procedures were performed in accordance with the relevant national guidelines for the Care and Use of Laboratory Animals (LANUV) and with approval from the authority for animal work at Heinrich Heine University Düsseldorf, Germany (Permit Number: O18/99).

Wild-type *Xenopus laevis* were obtained from the European Xenopus Resource Centre (EXRC) at University of Portsmouth, School of Biological Sciences, UK, or Xenopus 1, USA. Frog maintenance and care was conducted according to standard procedures in the AquaCore facility, University Freiburg, Medical Center (RI_00544) and based on recommendations provided by the international Xenopus community resource centers NXR (RRID:SCR_013731) and EXRC as well as by Xenbase (http://www.xenbase.org/, RRID:SCR_003280) ^63^. This work was done in compliance with German animal protection laws and was approved under Registrier-Nr. G-23/053 by the state of Baden-Württemberg.

### Mouse strains

We used *Rpgrip1l*^+/−^ mice for breeding and mated *Rpgrip1l*^+/−^ male and female mice to obtain wild-type and *Rpgrip1l*^−/−^ mouse embryos ^64^. *Rpgrip1l*^+/−^ mice were generated as follows: A *Rpgrip1l*-null allele was produced by using homologous recombination. The targeting construct was designed to introduce a PGKneo cassette replacing exon 4 and 5 of the *Rpgrip1l* coding sequence. The linearised vector was electroporated into R1 embryonic stem cells, and after G418 selection, all clones were screened by Southern blot analysis. Targeted embryonic stem cells were used to create chimeras that passed the *Rpgrip1l* mutation on to their progeny. F1 animals were intercrossed to derive homozygous mutant F2 newborns and embryos by timed mating. For genotyping, the following primers were used: KORpgrip1l_forward, 5′-GGCCTCCCCTTTGTCAT-3′; KOneo_forward, 5′-ACGAGTTCTTTCTGAGGGGATC-3′; and KORpgrip1l_reverse, 5′-CAGCTTTCCTTGTGTCTCTACTT-3′.

### Antibodies and plasmids

We used primary antibodies targeting acetylated α-tubulin (#sc-23950, Santa Cruz Biotechnology, Inc.), cleaved caspase-3 (#9661, Cell Signaling Technology), cleaved caspase-9 (#52873, Cell Signaling Technology), detyrosinated α-tubulin (#Ab01257-24.1, Absolute Antibody), FLAG (#200472, Agilent Technologies), MOAP1 (#PA5-112255, Thermo Fisher Scientific and #sc-271338, Santa Cruz Biotechnology, Inc.), MYC (#sc-789, Santa Cruz Biotechnology, Inc.), phospho-histone H3 (#06-570, Sigma-Aldrich) and γ-tubulin (#T6557, Sigma-Aldrich and #ab11317, Abcam). The generation of the polyclonal antibody against RPGRIP1L was described formerly ^64^. Appropriate anti-mouse, anti-rabbit and anti-goat antibodies conjugated with the fluorochromes DyLight 405, DyLight 488, Cy3, Alexa Fluor 405, Alexa Fluor 488, Alexa Fluor 647, FITC and CF568 were used as secondary antibodies. The following plasmids were used: IFT88-EYFP (kindly provided by David R. Beier) ^20,23^, FLAG-RPGRIP1L ^16^, FLAG-RPGRIP1L-T615P ^16^, FLAG-RID (RPGRIP1L) ^23^ and MYC-MOAP1. Mouse full-length MOAP1 was cloned into the MYC-tagged pCMV vector (Clontech).

### Cell culture, transfection and drug treatment

We used two different cell lines in this study. NIH3T3 cells (#ACC59) and HEK293 cells (#ACC35) were both purchased by the German Collection of Microorganisms and Cell Cultures GmbH (DSMZ). Cells were grown in DMEM containing 10% fetal calf serum (FCS), 1/100 (v/v) L-glutamine (Gibco), 1/100 (v/v) sodium pyruvate (Gibco), 1/100 (v/v) non-essential amino acids (Gibco) and 1/100 (v/v) pen/strep (Gibco) at 37°C and 5% CO_2_ until they reach confluency. Inactivation of mouse *Rpgrip1l*, *Cep290* and *Invs* in NIH3T3 cells and human *RPGRIP1L* in HEK293 cells was performed as previously described ^20,21^. The following clones were used: *Rpgrip1l*^-/-^ NIH3T3 cells (clone 10-61) ^21^, *Cep290*^-/-^ NIH3T3 cells (clone 39-51)^20^, *Invs*^-/-^ NIH3T3 cells (clones 48-20)^20^ and *RPGRIP1L*^-/-^ HEK293 cells (clone 1-7)^20^. Inactivation of mouse *Tctn1* in NIH3T3 cells was carried out as described below in the part “CRISPR/Cas9-mediated gene inactivation”.

Ciliogenesis was induced by serum starvation for at least 24 hours. Cells were treated with 20 µM MG132 (#BML-PI102-0025; Enzo Life Sciences) for 6 h, with 5 µM SFN (#ALX-350-230-M010; Enzo Life Sciences) for 20 h, with 10 nM RAP (#9904S; Cell Signaling Technology) for 20 h or with 20 µM EUP (#SML1689-5MG; Merck) for 24 h. As a solvent control, cells were treated with DMSO for the same time, respectively. The transfection of NIH3T3 and HEK293 cells with the respective constructs and siRNAs used in different experiments was carried out by using Lipofectamine 3000 (Thermo Fisher Scientific) according to the manufacturer’s manual.

### Coimmunoprecipitation

Fusion proteins, MYC-tagged MOAP1 full-length protein and the RID domain of RPGRIP1L with a FLAG-tag, were overexpressed in NIH3T3 cells for 24 h. Afterwards, cells were lysed on ice in lysis buffer (50 mM Tris-HCl, pH 7.4, 150 mM NaCl, 1% NP-40, and 0.25% Na-deoxycholate) supplemented with complete protease inhibitor cocktail (#11836153001, Roche) and PMSF (#93482, Sigma-Aldrich). Lysates were firstly precleared with protein G–agarose (#16-266, EMD Millipore) overnight and then incubated with anti-FLAG or anti-MYC protein G–agarose overnight. Subsequently, beads with bound protein complexes were washed in lysis buffer, put into 4× sample buffer (50 mM Tris-HCl, pH 6.8, glycerol, 2% SDS, 20% β-mercaptoethanol and bromophenol blue), heated for 10 min at 95°C and precipitated by centrifugation. The supernatant was separated by using SDS-PAGE and then immunoblotted.

### Drug treatment of *Xenopus* embryos

Injected and control embryos were cultured until stage 13 was reached. To permeabilise the fertilization membrane, embryos were incubated in 1/3x MR containing 8µg/ml proteinase K (Sigma Aldrich, #P6556-500MG) for 3 minutes at room temperature (RT), then rinsed twice in 1/3x MR and immediately transferred into 12 well plates containing 1/3x MR. Treatment with 20 µM Sulforaphane (Enzo Life sciences, #ALX-350-230-M010) in DMSO (Honeywell, #472301-500ML) or vehicle treatment (0.2-0.5% DMSO) was conducted at 16°C with daily media changes and protected from light until stage 32-34 was reached.

### Fertilization and microinjection of *Xenopus* embryos

Wild-type *X. laevis* eggs were collected and *in-vitro* fertilized using wild-type sperm. Embryos were then microinjected using standard procedures ^65^. Injections were conducted at two to eight cell stage in a solution of 1/3x MR (Modified Frog Ringer’s solution pH 7.6, 33 mM NaCl, 6 mM KCl, 0.66 mM CaCl2, 0.33 mM MgCl2, 1.66 mM HEPES) with 2.5% Ficoll (PM 500, GE Healthcare, #17-0300-50) using a PicoSpritzer 3000 setup targeting the ectoderm (animal hemisphere injections). Embryos were injected with morpholino oligonucleotides (MOs, Gene tools) and membrane-GFP mRNA (50 ng/μl, gift from the Harland Lab) as a lineage tracer. The droplet size was calibrated to 7-8 nl. After injection, embryos were transferred to a solution of 1/3x MR containing 50 µg/ml Gentamycin.

MO sequences:

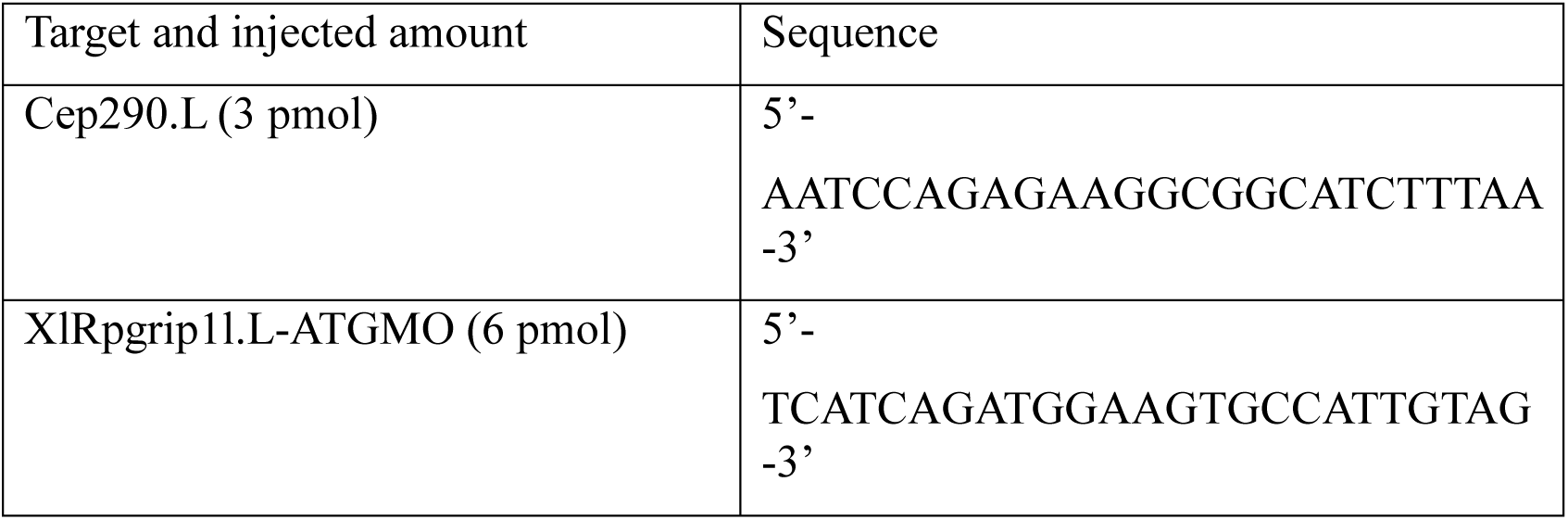

### High-resolution imaging, super-resolution imaging and 3D reconstruction

The acquisition of high-resolution images was performed by using an Olympus FluoView FV3000 confocal laser scanning microscope equipped with TruSpectral detectors and 405 nm, 488 nm, 561 nm and 640 nm lasers.

Super-resolution images were acquired with a ZEISS LSM 980 Airyscan2 with a 63x oil immersion objective NA 1.4. The super-resolution mode of the Airyscan2 module was used and raw data was further processed with the Hyugens software. 3D reconstruction, image and movie recording was performed in the ARIVIS PRO software.

Single-molecule localisation microscopy (SMLM) was accomplished by using dSTORM. dSTORM data were obtained with a Bruker Vutara VXL super-resolution microscope equipped with an ORCA-Fusion BT sCMOS camera, with 405 nm, 488 nm and 555 nm lasers and with the bi-plane detection technique to achieve 3D sub-diffraction resolution. Image processing and movie recording was performed in the SRX software.

### Immunofluorescence

For immunofluorescence on NIH3T3 cells and HEK293 cells, cells were plated on coverslips until they reach confluency. Cells were serum-starved for 24 hours and fixed with 4% PFA for 1 hour at 4°C or with 100% MeOH for 5 minutes at -20°C. Fixed cells were rinsed three times with PBS, followed by a permeabilisation step with PBS/0.5% Triton-X-100 for 10 minutes. The samples were washed three times in PBST (PBS/0.1% Triton-X-100) and blocking was performed by incubation in PBST containing 10% donkey serum for at least 1 hour at room temperature. Diluted primary antibodies (in PBS/0.1% Triton-X-100) were incubated overnight at 4°C. After three washing steps with PBST, incubation with fluorescent secondary antibody (in PBS/0.1% Triton-X-100) was performed at room temperature for 1 hour followed by several washing steps and subsequent embedding with Mowiol optionally containing DAPI (#1.24653; Merck).

For immunofluorescence on murine limb buds, mouse embryos at E12.5 were isolated and limb buds were dissected. Limb buds were fixed in 4% PFA for 1 hour and incubated in 30% sucrose (in PBS) overnight at 4°C. The next day, limb buds were embedded in Tissue-Tec O.C.T. compound (#4583; Sakura Finetechnical) and freezed at -80°C. Transverse cryostat sections (6 µm) were prepared. Limb bud sections were rinsed in PBS, permeabilised with PBS/0.5% Triton X-100 for 10 minutes and washed three times in PBS/0.1% Triton X-100. Blocking was performed with 10% donkey serum in PBST. Subsequently the sections were incubated with primary antibodies diluted in PBS/0.1% Triton X-100 overnight at 4°C. On the next day, the sections were washed three times with PBS/0.1% Triton X-100 and incubated with the secondary antibodies diluted in PBS/0.1% Triton X-100 for 2 hours. After several washing steps the sections were embedded in Mowiol containing DAPI (#1.24653; Merck).

### *In situ* PLA

To test whether RPGRIP1L and MOAP1 are in close proximity (< 40 nm), we used the *in situ* proximity ligation assay (PLA or DUOLINK^TM^). *In situ* PLA was performed as described before ^26^ according to the manufacturer’s manual (OLinkBioscience). RPGRIP1L and MOAP1 were recognised by specific primary antibodies. The cilium in the *in situ* PLA experiments was marked by the product of a transiently transfected IFT88-EYFP construct (Fig. 5b).

### Statistical analysis

Data are presented as mean ± s.e.m. or as median with quartiles. Two-tailed t test with Welch’s correction was performed for all data in which two normally distributed datasets were compared. Ordinary one-way analysis of variance (ANOVA) with Tukey’s multiple comparison test was used for all data in which more than two normally distributed datasets were compared. Mann-Whitney U test was performed for datasets that were not normally distributed. Kruskal-Wallis with Dunn’s multiple comparison test was used for comparison of more than two datasets with small or not normally distributed data. Brown-Forsythe and Welch ANOVA test with Dunnett’s T3 multiple comparisons test was used for all data in which three or more normally distributed datasets were compared without assuming that the datasets have equal variances. A P-value <0.05 was considered to be statistically significant (one asterisk), a P-value <0.01 was defined as highly significant (two asterisks), a P-value <0.001 was noted as very highly significant (three asterisks) and a P-value <0.0001 was considered to be extremely significant (four asterisks). All statistical data analysis and graph illustrations were performed using GraphPad Prism (GraphPad Software).

### Fluorescence-based quantifications of ciliary protein amounts

The amount of ciliary proteins was determined by the comparative measurement of fluorescence intensity. Intensity of ciliary protein staining on z-stacks was quantified by using ImageJ (National Institutes of Health). In principle, intensity of ciliary proteins based on immunofluorescence staining was measured as described before ^21–23,33,66–68^. To improve the accuracy of the method, we quantified the fluorescence signal of the ciliary protein of interest in five optical sections of each z-stack. For the quantification of axonemal TRAIL-R1, we used the area marked by detyrosinated α-tubulin and quantified the average pixel intensity to take the cilia length into account thereby making the data of wild-type and *Rpgrip1l*^-/-^ cilia comparable. For the quantification of MOAP1, TRAIL-R1 and TNF-R1 at the basal body and the daughter centriole, we selected the regions of interest by encircling the respective areas labelled by γ-tubulin and measured the total pixel intensity. The labelling of the selected region was outlined with the freehand selection tool and the mean intensity of the desired channel inside the area measured in an 8-bit scale (0-255). To get rid of the ratio of unspecific (background) staining, we subtracted the mean value of the average pixel intensity (in the case of axonemal TRAIL-R1) or of the total pixel intensity (for MOAP1, TRAIL-R1 and TNF-R1 at the basal body and the daughter centriole) of three neighbouring regions free from specific staining. The quantified areas of the specific staining and the unspecific staining of every individual measurement have to be equal in size.

### Quantifications of cells undergoing PCD

The number of cells undergoing PCD was determined via cell counting. Total cell number was ascertained by counting cell nuclei marked by DAPI. Cells undergoing PCD were counted via CC3- or CC9-positive signals and related to the total cell number. Finally, the percentage of cells undergoing PCD was normalised to the (untreated) wild-type status.

### RNAi experiments

200 pmol of a Silencer® Select siRNA against *Moap1* (#4392420, Thermo Fisher Scientific) was transfected into *Rpgrip1l*^-/-^ NIH3T3 cells. Scrambled siRNA was used as a negative control.

### TUNEL assay in *Xenopus* embryos

Embryos were fixed in 1x MEMFA (100 mM MOPS pH7.4, 2 mM EGTA, 1 mM MgSO4, 3.7% (v/v) Formaldehyde) overnight at 4°C or for 2 hours at RT, and stored in 100% Ethanol at - 20°C until use. Embryos were bleached before staining ^65^. TUNEL staining was performed as described in ^69^ using Terminal Deoxynucleotidyl Transferase Kit (Invitrogen #10533065), dig-UTP (Roche, #3359247910), anti-Digoxigenin AP Fab fragments (Roche, #11093274910) and NBT/BCIP (Roche, #11681451001). Staining was stopped with 100% Methanol (Roth, #8388.2), samples were then fixed briefly with 4% PFA (Roth, #0335.1) in PBS (Phosphate buffered saline, 10 mM Na2HPO4, 1.8 mM KH2PO4, 137 mM NaCl, 2.7 mM KCl) and imaged on a Zeiss Stemi508 with Axiocam208-color, and images were adjusted for colour balance, brightness and contrast using Adobe Photoshop.

### Western Blotting

Western blot studies were performed as described earlier ^20^. We used an anti-MOAP1 antibody (#sc-271338, Santa Cruz Biotechnology, Inc.). An anti-HPRT antibody (#A2066, Sigma-Aldrich) was used to detect the loading control. Protein bands were visualised by the LAS-4000 mini (Fujifilm) and band intensities were measured by using ImageJ (National Institutes of Health).

### Yeast two-hybrid screen

A GAL4-based yeast two-hybrid system (BD) and a mouse E11.5 cDNA library (BD) were used in the yeast two-hybrid screen. The yeast strain AH109 was used as host carrying the *HIS3* (histidine), *ADE2* (adenine), *MEL1* (α-galactosidase) and *LacZ* (β-galactosidase) reporter genes. Interactions were investigated by evaluation of reporter gene activation via growth on selective media (*HIS3* and *ADE2* reporter genes), α-galactosidase colorimetric plate assays (*MEL1* reporter gene), and β-galactosidase colorimetric filter lift assays (*LacZ* reporter gene). The evaluation of the suspected protein interactions was conducted using growth and selective media, along with staining in α- und β-galactosidase activity assays.

## Acknowledgements

We express our gratitude to Ilona Renken-Olthoff and the entire team of the Health and Medical University for their support. The authors greatly acknowledge France Lam of the IBPS Imaging Facility for her help in high resolution imaging and 3D image processing. Our special thanks go to the Bruker Corporation, especially to Corentin Rousset and Clemens Schneider, for the opportunity to test the Vutara VXL super-resolution microscope and to acquire dSTORM data. Furthermore, we are grateful to David R. Beier for providing the IFT88-EYFP construct. We thank Henrike Tietz for critical reading of the manuscript and Johanna Maria Lier for generating parts of schematic illustrations. This work was supported by the Deutsche Forschungsgemeinschaft (Sonderforschungsbereiche 590 and 612; U.R.). The IBPS Imaging facility is supported by Region-Île-de-France, Sorbonne-University and CNRS.

## Author contributions

Conceptualisation: C.G. Investigation: A.W., S.L.H., S.K., M.B., C.O., P.W. and C.G. Visualisation: A.W., S.L.H., S.K., C.O. and C.G. Provision of resources: S.S. Funding acquisition: U.R. Project administration: C.G. Supervision: T.G., U.R., S.S.M., P.W., T.P. and C.G. Writing – original draft: C.G.

## Expanded View data

**Fig. EV1:**
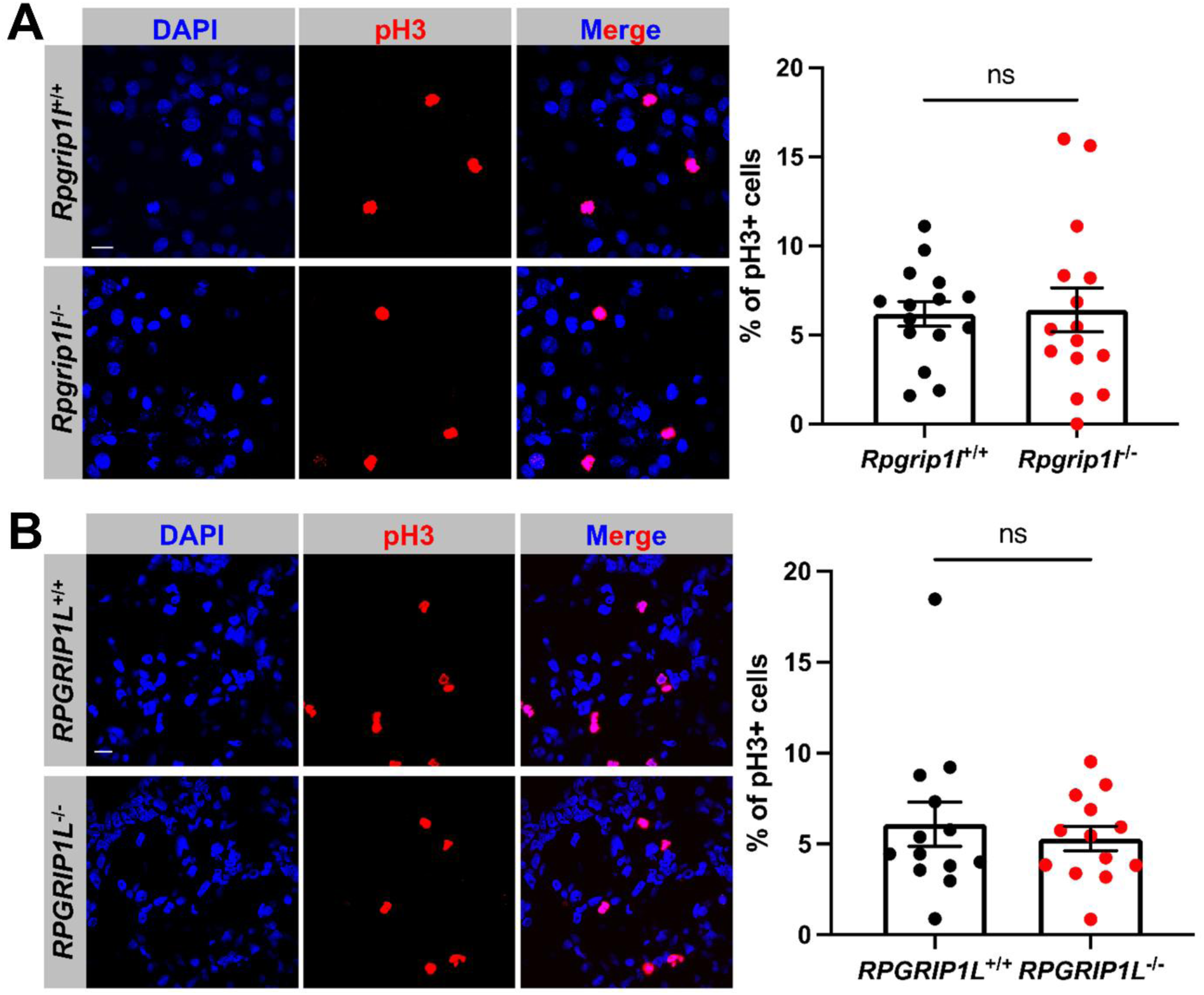
Loss of RPGRIP1L does not alter proliferation in NIH3T3 and HEK293 cells. (**A**) Immunofluorescence on NIH3T3 cells. Cell nuclei are stained in blue by DAPI. Proliferating cells are marked in red by pH3. The scale bar (in white) represents a length of 20 μm. At least 840 cells per genotype were used for these measurements, pooled from three replicates. Data shown as mean ± s.e.m. Statistical evaluation was performed by using two-tailed unpaired t test with Welch’s correction. ns = not significant. (**B**) Immunofluorescence on HEK293 cells. Cell nuclei are stained in blue by DAPI. Proliferating cells are marked in red by pH3. The scale bar (in white) represents a length of 20 μm. At least 1600 cells per genotype were used for quantification, pooled from three replicates. Data shown as mean ± s.e.m. Statistical evaluation was performed by using two-tailed unpaired t test with Welch’s correction. ns = not significant.

**Fig. EV2:**
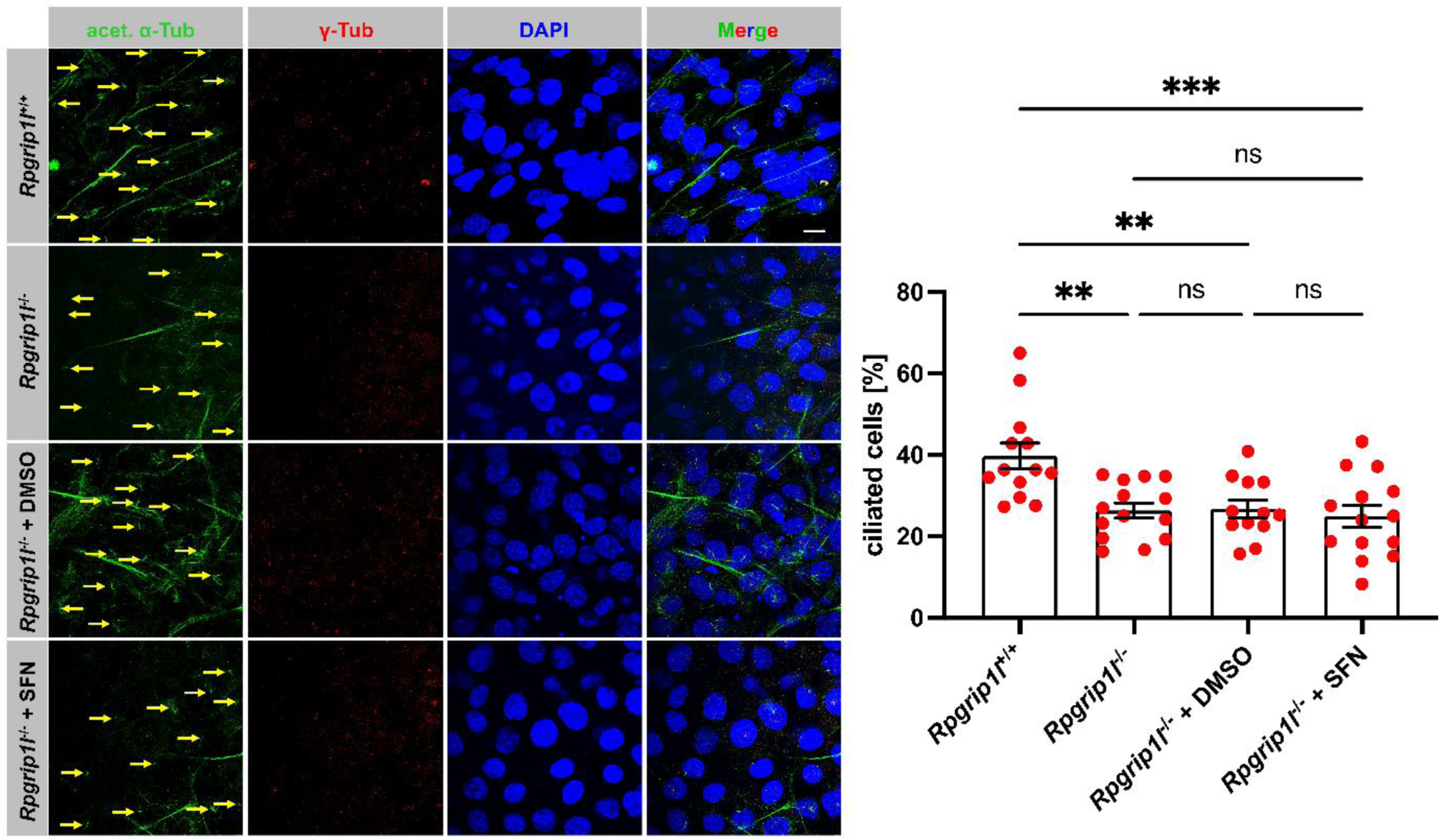
The number of ciliated NIH3T3 cells is decreased in the absence of RPGRIP1L and the treatment of *Rpgrip1l*^-/-^ NIH3T3 cells with SFN does not rescue the number of ciliated cells. Immunofluorescence on serum-starved NIH3T3 cells. Cell nuclei are stained in blue by DAPI. The ciliary axoneme is stained in green by acetylated α-tubulin and the BB in blue by γ-tubulin. Yellow arrows point to cilia. The scale bar (in white) represents a length of 10 μm. At least 470 cells per genotype and treatment were used for quantification, pooled from three replicates. Data shown as mean ± s.e.m. Statistical evaluation was performed by using ordinary one-way ANOVA with Tukey’s multiple comparisons test. Asterisks denote statistical significance (*Rpgrip1l*^+/+^ vs. *Rpgrip1l*^-/-^: ** *P* = 0.0023; *Rpgrip1l*^+/+^ vs. *Rpgrip1l*^-/-^ + DMSO: ** *P* = 0.0049; *Rpgrip1l*^+/+^ vs. *Rpgrip1l*^-/-^ + SFN: *** *P* = 0.0006). ns = not significant.

**Fig. EV3:**
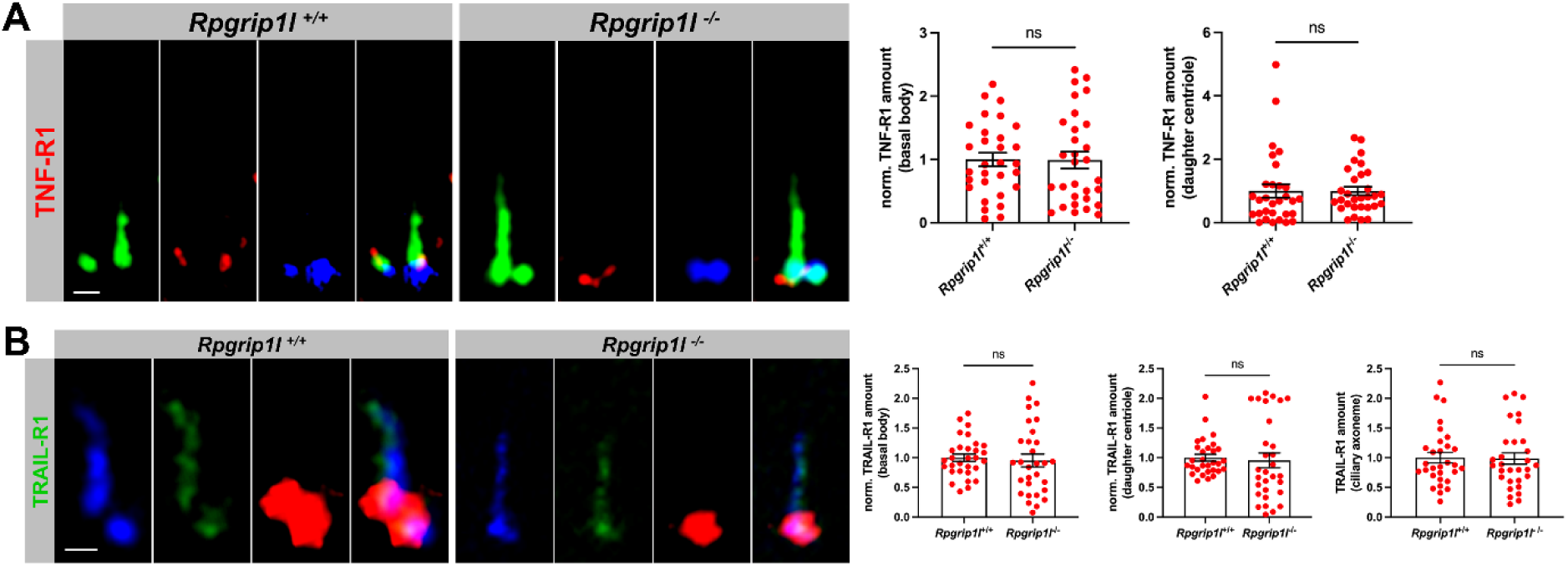
The death receptors TNF-R1 and TRAIL-R1 are located at cilia and their ciliary localisation is independent of RPGRIP1L. (**A, B**) Immunofluorescence on serum-starved NIH3T3 cells. (**A**) The ciliary axoneme is stained in green by detyrosinated α-tubulin and the BB in blue by γ-tubulin. The scale bar (in white) represents a length of 1 μm. 30 cilia per genotype were used for quantification, pooled from three replicates. Data shown as mean ± s.e.m. Statistical evaluation was performed by using two-tailed unpaired t test with Welch’s correction (quantification at basal body) and by using two-tailed Mann-Whitney U test (quantification at daughter centriole). ns = not significant. (**B**) The ciliary axoneme is stained in blue by detyrosinated α-tubulin and the BB in red by γ-tubulin. The scale bar (in white) represents a length of 1 μm. 30 cilia per genotype were used for quantification, pooled from three replicates. Data shown as mean ± s.e.m. Statistical evaluation was performed by using two-tailed unpaired t test with Welch’s correction (quantification at basal body), by using two-tailed Mann-Whitney U test (quantification at daughter centriole) and by using two-tailed Mann-Whitney U test (quantification at ciliary axoneme). ns = not significant.

**Fig. EV4:**
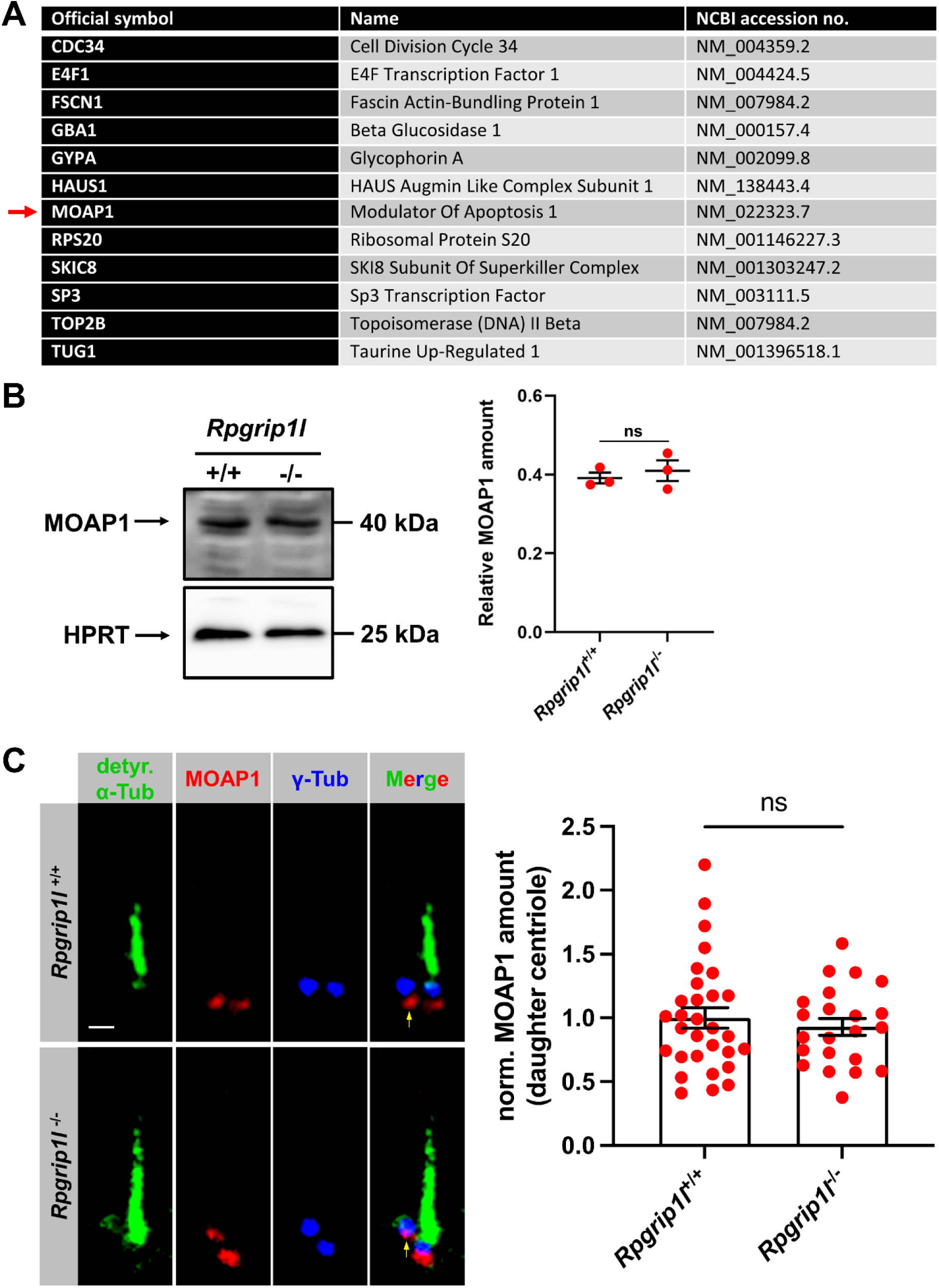
MOAP1 was identified as an interaction partner of RPGRIP1L and its overall cellular amount as well as its amount at the daughter centriole is unaltered in the absence of RPGRIP1L. (**A**) List of RPGRIP1L-interacting proteins identified via yeast two-hybrid screen. Accession numbers were obtained from GenBank. Red arrow points to the most promising interaction partner in regard to PCD – MOAP1. (**B**) Western blot analysis with lysates extracted from *Rpgrip1l*^+/+^ and *Rpgrip1l*^-/-^ NIH3T3 cells. HPRT serves as loading control. (**C**) Immunofluorescence on serum-starved NIH3T3 cells. The ciliary axoneme is stained in green by detyrosinated α-tubulin and the BB in blue by γ-tubulin. The scale bar (in white) represents a length of 1 μm. 30 cilia (*Rpgrip1l*^+/+^) and 22 cilia (*Rpgrip1l*^-/-^) were used for quantification, pooled from three replicates. Data shown as mean ± s.e.m. Statistical evaluation was performed by using two-tailed unpaired t test with Welch’s correction. ns = not significant.

**Fig. EV5:**
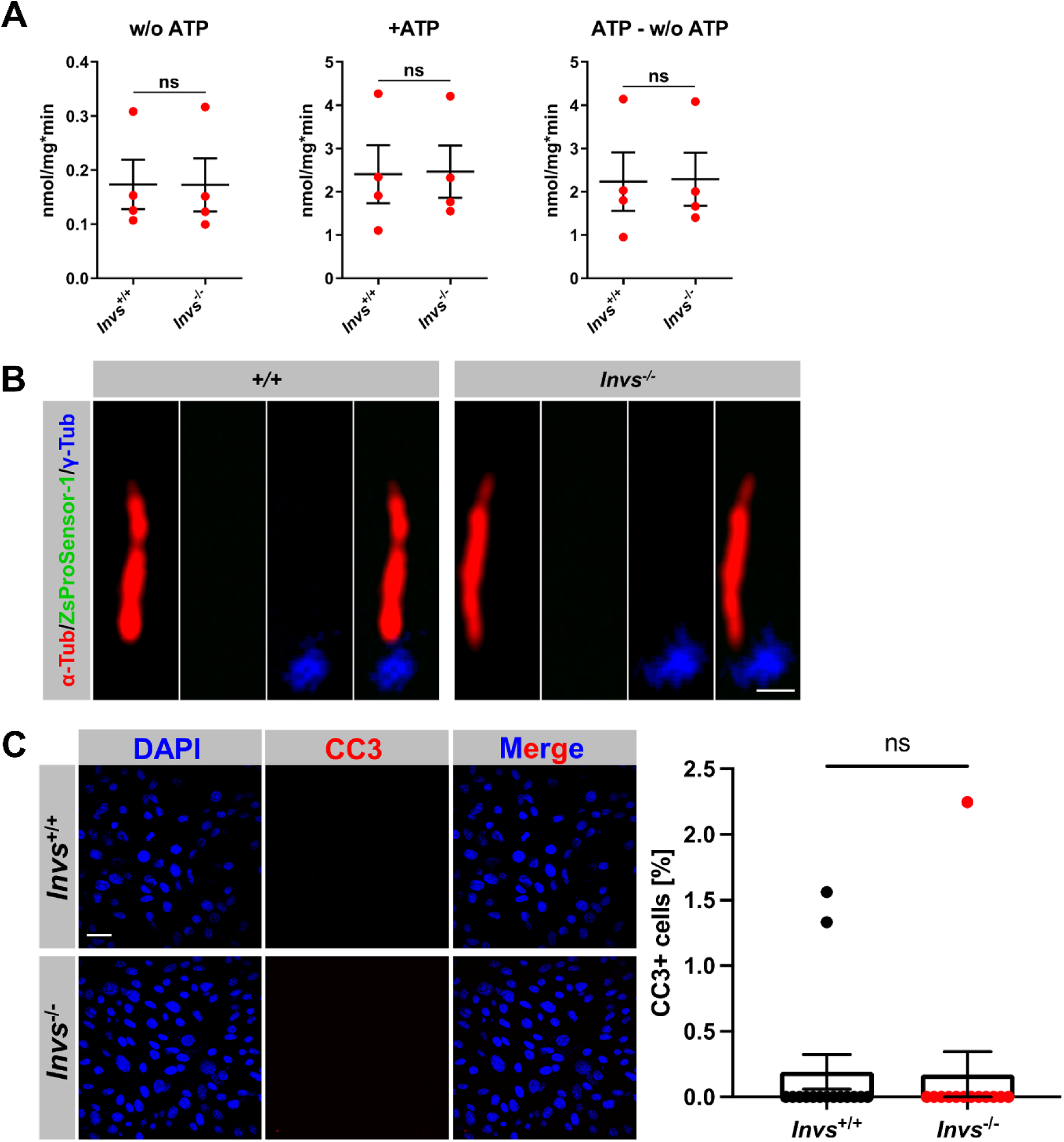
Proteasomal activity and PCD are unaltered in *Inv*^-/-^ NIH3T3 cells. (**A**) Proteasomal activity was quantified in total cell lysates of serum-starved NIH3T3 cells. The activity of the 20S proteasomal subunit was measured without ATP. The activity of the 20S proteasomal subunit and the 26S proteasome together was quantified after ATP addition. The activity of the 26S proteasome was determined by subtracting the value measured for the activity of the 20S proteasomal subunit from the value measured for the activity of the 20S proteasomal subunit and the 26S proteasome together. ns = not significant. (**B**) Fluorescence-based proteasome activity assay on serum-starved NIH3T3 cells. The ciliary axoneme is stained in green by acetylated α-tubulin and the BB in blue by γ-tubulin. The ZsProSensor-1 protein signal shines green. The scale bar (in white) represents a length of 0.5 μm. (**C**) Immunofluorescence on serum-starved NIH3T3 cells. Cell nuclei are stained in blue by DAPI. Cells undergoing PCD are marked in red by CC3. The scale bar (in white) represents a length of 20 μm. At least 980 cells per genotype were used for these measurements, pooled from three replicates. Data shown as mean ± s.e.m. Statistical evaluation was performed by using two-tailed Mann-Whitney U test. ns = not significant.

